# MicroRNAs are required within a critical time window to define neural patterning during early human brain development

**DOI:** 10.64898/2026.03.30.715268

**Authors:** Lisa Emmenegger, Cledi Alicia Cerda Jara, Matilde Ercolano, Jaden Loebert, Nicolas Morando, Poojashree Bhaskar, Ivano Legnini, Agnieszka Rybak-Wolf, Nikolaus Rajewsky

**Affiliations:** Laboratory for Systems Biology of Gene Regulatory Elements, Berlin Institute for Medical Systems Biology (BIMSB), Max Delbrück Center for Molecular Medicine (MDC) in the Helmholtz Association, Hannoversche Str. 28, 10115 Berlin, Germany; Organoid Platform, Berlin Institute for Medical Systems Biology, Max Delbrück Center for Molecular Medicine in the Helmholtz Association (MDC), Berlin, Germany; Instituto de Investigaciones Biomédicas en Retrovirus y Sida (INBIRS), Consejo Nacional de Investigaciones Científicas y Técnicas (CONICET)-Universidad de Buenos Aires, Buenos Aires 1121, Argentina; Human Technopole, Viale Rita Levi-Montalcini 1, 20157 Milano, Italy; Charité - Universitätsmedizin, Charitéplatz 1, 10117 Berlin, Germany; German Center for Cardiovascular Research (DZHK), Berlin, Germany; NeuroCure Cluster of Excellence, Berlin, Germany; National Center for Tumor Diseases (NCT), Berlin, Germany; ImmunoPreCept, Cluster of Excellence, Berlin, Germany; Einstein Center for Early Disease Interception, Berlin, Germany

## Abstract

MicroRNAs (miRNAs) are key post-transcriptional regulators of cell state transitions, yet their function in early human brain development is largely unknown. Here, we present a longitudinal analysis of miRNA function in developing human forebrain organoids. We show that mRNAs and miRNAs expression mirrors known developmental gene programs and that miRNA biogenesis peaks at neural commitment. To test the function of miRNAs in regulating commitment, we impaired their biogenesis at defined stages. miRNA disruption during pre-neuronal commitment caused severe patterning defects, whereas post-commitment perturbation had minimal impact on forebrain identity. We show that miRNA loss during pre-commitment increased WNT and BMP signaling, thus shifting cell fates towards non-forebrain identity such as midbrain/hindbrain. These effects could be partially rescued by expressing five miRNAs. Our findings uncover a critical time window where miRNAs regulate morphogen signaling in early human neurodevelopment, establishing them as essential temporal determinants of cell fate and brain regional identity.

## Introduction

Transcriptional as well as post-transcriptional mechanisms orchestrate spatial and temporal gene expression programs that control neurogenesis and tissue patterning in animals (Gibney and Nolan, 2010; Spitz and Furlong, 2012; Rajman and Schratt, 2017). MiRNAs are conserved, small non-coding RNAs (20-24 nt) that regulate post-transcriptional gene expression by targeting binding sites in 3’ untranslated regions (3’ UTRs) of mRNAs (reviewed in Bartel, 2018; Rajewsky 2006; Aigner et al., 2026). After transcription of a miRNA gene by RNA Polymerase II, and subsequent processing steps by DROSHA and DICER, miRNAs guide the RNA-induced silencing complex (RISC) to target mRNAs, typically leading to degradation, translational repression, and regulation of mRNA localization (Bartel, 2018; Ebert and Sharp, 2012; Mendonsa et al., 2023). Constitutive depletion of core components of the miRNA machinery is embryonically lethal (Dicer: Bernstein et al., 2003; Ago2: Morita et al., 2007; DGCR8: Wang et al., 2007, reviewed in detail by Alberti and Cochella, 2017), indicating essential roles of miRNAs in early development.

In fact, miRNAs were originally discovered as heterochronic genes that regulate the timing of developmental transitions by repressing stage-specific cell fate determinants (Ruvkun et al., 1989; Lee et al., 1993; Wightman et al., 1993). Studies in many different species further support this conserved temporal regulatory role (D. rerio: Wienholds et al., 2003, and Giraldez et al., 2005; C. elegans: Stoeckius et al., 2009; S. purpuratus: Song et al., 2012; Drosophila: Kataoka et al., 2001). For example, in sea urchins, loss of Drosha disrupts miRNA biogenesis and leads to defective gastrulation (Song et al., 2012). Furthermore, canonical miRNAs are linked to the emergence of multicellularity (Mattick, 2004; Dexheimer and Cochella, 2020). Moreover, the evolution of miRNA genes, their expression and targeting, has been strongly associated with the evolution of complex nervous systems (Chen and Rajewsky 2007; Heimberg et al., 2008; Fromm et al., 2015; Zolotarov et al., 2022). However, little is known about the role of miRNAs in the developing human brain.

Here, we set out to study the function of miRNAs in early human brain development. We leveraged the power human neural organoids, which have recently emerged as a powerful model to study early human neurodevelopment under both physiological and pathological conditions. These 3D models mirror important aspects of tissue architecture, cellular complexity, and developmental timing (Eiraku et al., 2008; Lancaster et al., 2013; Lancaster and Knoblich, 2014; Amin et al., 2024). Importantly, the embryonic development of specific brain regions in organoids can be consistently modelled by the addition of extrinsic patterning factors, which modulate morphogen signaling (Paşca et al., 2015; Qian et al., 2018). For instance, morphogens like WNT and BMP define distinct cell identities along the anterior-posterior axis of the developing vertebrate neural tube (Pires-daSilva and Sommer, 2003).

We employed a well-established forebrain patterning protocol (Walsh et al., 2024) across two independent iPSC lines. We performed almost all experiments in these two independent lines to ensure reproducibility across different genetic backgrounds. We first show that miRNA and mRNA expression in these systems faithfully reflect what is known about human fetal RNA expression and developmental programs. However, we found that both miRNAs and their biogenesis machinery undergo dynamic changes throughout human forebrain development. In particular, protein expression of crucial components of miRNA biogenesis and function (DROSHA, DICER, AGO2) strongly peaked at the stage of neuronal commitment. By perturbing miRNA biogenesis through overexpression of a dominant-negative DROSHA mutant (p.E1147K; Rakheja et al., 2014; Torrezan et al., 2014), which is known to specifically impair maturation of miRNAs, we uncovered a critical developmental window during which miRNAs orchestrate, post-transcriptionally, forebrain specification. We identified a mechanistic link between the temporal requirement of miRNAs and the direct regulation of WNT/BMP pathways. This miRNA-morphogen regulation provides a mechanistic explanation for the observed cell fate switch. Moreover, the gain of function of forebrain-enriched miRNAs, before cell commitment, partially restores the regionalization defects. Overall, our results reveal that miRNAs exert a temporally defined control over human neural lineage progression and cortical organization, by constraining WNT/BMP signaling, at the earliest pre-commitment stages of development. Our results show that miRNA are essential regulators of early human brain patterning. Moreover, our findings show that miRNAs can be deployed to alter the patterning of brain organoids, adding them to the toolbox for organoid engineering.

## Methods

### 1. Human induced pluripotent stem cell lines

In this study we used two independent human induced pluripotent stem cell (hiPSC) lines, derived from different donors: 1. Gibco™ Episomal hiPSC Line catalog no. A18945, from Thermo Fisher Scientific and 2. Tissue-T106 hiPSC line). Cells were cultured under hypoxic conditions (5% O_2_) at 37°C in E8 Flex medium with supplement (Thermo Fisher, catalog no. A2858501). They were passaged every 3-4 days using TrypleE and seeded on Geltrex-coated plates (Gibco catalog). no. A1413302). At the time of splitting, cells were kept in E8 Flex medium supplemented with 10 μM Rho-associated protein kinase (ROCK) inhibitor to promote cell adhesion. Within 24 hours, the medium was replaced with E8 Flex without ROCK inhibitor. Cells were routinely tested for mycoplasma contamination.

### 2. Cloning and genetic engineering

The original plasmid containing the dominant-negative *DROSHA* variant (“V5-Drosha E1147K-pcDNA”) was kindly provided by the groups of Joshua Mendell and Kenneth Chen. This mutation affects the catalytic domain of DROSHA and functions through a dominant-negative mechanism, interfering with wild-type *DROSHA* (Rakheja et al., 2014; Torrezan et al., 2014). The donor plasmid, “e.PB_PuroR_NeonGreen_TRE,” is a PiggyBac vector, which contains a doxycycline-inducible promoter with a NeonGreen cassette directly downstream. For simplicity, in the whole manuscript we refer to NeonGreen as “GFP”. This plasmid was digested with appropriate restriction enzymes (Fast Digest, Thermo Scientific) and the resulting products were purified from a 1% agarose gel using a Gel DNA recovery kit (Zymoclean, Zymo catalog no. D4001). The *DROSHA* insert was amplified by PCR from “V5-DROSHA E1147K-pcDNA”, resolved on a 1% agarose gel, purified, and subsequently subcloned into the digested PiggyBac vector. Vector and insert were assembled using the HiFi DNA Assembly Master Mix (NEB, catalog no. E2621). Cloning was performed via transformation of chemically competent *E. coli* bacteria (Mix&Go Competent Cells DH5α, Zymo catalog no. T3007), with ampicillin used for selection. Final plasmids were isolated using the ZymoPURE Plasmid Miniprep kit (Zymo catalog no. D4214), and the correct sequences were confirmed by Sanger sequencing and by whole plasmid sequencing. HiPSC lines were next genetically engineered to genomically integrate the doxycycline-inducible system (Tet-On) to overexpress the *DROSHA* dominant-negative mutant (c. G3439A, p. E114K) (Rakheja et al., 2014; Torrezan et al., 2014). The tetracycline-responsive element (TRE) of the Tet-On system is fused to the hUBC (human ubiquitin C) promoter. This promoter was reported to drive transgene expression more extensively in early neurons, suggesting, in our case study, a predominant impact of *DROSHA* perturbation on neuronal populations (Wilhelm et al., 2011). Stable hiPSC lines were generated by seeding 400,000 hiPSCs into a Geltrex-coated well of a 12-well plate and transfecting them using Lipofectamine Stem (Thermo Fisher, catalog no. STEM00001) diluted in OPTI-MEM (Thermo Fisher, catalog no. 31985062). The transfection mix contained 600 ng of PiggyBac vector and 300 ng of a plasmid encoding a hyperactive PiggyBac transposase. To monitor transfection efficiency, a plasmid encoding constitutive GFP was included as a control (pmax-GFP, Lonza, catalog no. V4YP-1A24). The medium was replaced with E8 Flex the morning after transfection. Following 4-5 days of recovery, transfected cells were subjected to antibiotic selection with 1 ug/ml of puromycin for 10 days. Only cells that had stably integrated the construct into their genome survived the selection process, yielding multiclonal populations, which likely harbored varying copy numbers of the transgene. Upon doxycycline treatment, *GFP* and *DROSHA* mutants are transcribed as one transcriptional unit, and the presence of a T2A self-cleaving peptide in between ensures the translation into independent proteins.

### 3. Human forebrain organoids generation

We generated hiPSC-derived forebrain organoids according to an optimized version of a previously described protocol (Walsh et al., 2024). On day 0, hiPSCs were dissociated to single cells using TrypleE, and 9,000 cells per well were seeded into 96-well plates to form embryoid bodies (EBs) in 100 µl of hiPSC culture medium, supplemented with 50 µM ROCK inhibitor and 5 µM WNT inhibitor (XAV939 5 µM Enzo Life Sciences, catalog no. 15137408, hereafter XAV). On the next day, half of the medium was replaced with E6 (Thermo Fisher catalog no. A1516401) supplemented with 1% penicillin-streptomycin, 5 µM XAV, 10 µM TGF-β pathway inhibitor (SB431542, Miltenyi Biotec, catalog no. 130-106-275, herein SB), and 0.1 µM BMP inhibitor (LDN193189, Miltenyi Biotec, catalog no. 130-106-540, herein LDN). The medium was replaced every other day until day 5, after which the neural induction was continued without XAV supplementation, and EBs were cultured at 37 °C in a tissue culture incubator with 5% CO₂ and 20% O_2_. On day 9, the medium was switched to neural differentiation medium (COM1), composed of a 1:1 mixture of Dulbecco’s modified Eagle medium (DMEM)/F12 (Thermo Fisher, catalog no. 11320033) and Neurobasal Plus (Thermo Fisher, catalog no. A3582901), 1x GlutaMAX (Thermo Fisher, catalog no. 35050038), 1x Minimal Essential Medium-Non Essential Amino Acid (Sigma Aldrich catalog no. M7145, hereon MEM-NEAA), 1x N2 supplement (STEMCELL Technologies, catalog no. 07152), 1x B27 without vitamin A (STEMCELL Technologies, catalog no. 05731), 2.5ug/ml insulin (Sigma Aldrich, catalog no. 19278), 0.05 mM 2-mercaptoethanol (Sigma Aldrich, catalog no. 805740), 200 nM ascorbic acid (Sigma Aldrich, catalog no. 95209), 1x antibiotic-antimycotic (Sigma Aldrich, catalog no. 15240062). COM1 was further supplemented with 10 ng/mL leukemia inhibitory factor (LIF), a STAT3 pathway activator, to promote outer radial glia (oRG) proliferation and differentiation (Walsh et al., 2024). On days 10-11, organoids underwent liquid embedding in cold COM1 medium supplemented with 2% Geltrex and 10 ng/mL LIF. Organoids were then transferred to six-well plates and cultured on an orbital shaker (80 rpm) to facilitate the formation of 3D structures. From days 16-17 onward, organoids were maintained in neural maturation medium (COM2), composed of a 1:1 mixture of DMEM/F12 and Neurobasal Plus, N2 supplement, B27 with vitamin A (Stem Cell Technologies, catalog no. 05711), 2.5 μg/ml insulin, 0.05 mM 2-mercaptoethanol, 1x GlutaMAX, 1x MEM-NEAA and 200 nM ascorbic acid. Media changes were performed every 3 days. Live organoid imaging was conducted using NIKON Eclipse Ts2 microscope. Experiments presented in **Figures 1-2** and **Supplementaries 1-4** were conducted on uninduced, genetically engineered organoids, integrating the doxycycline-inducible system (described in Section 4 of Methods). These organoids served as isogenic controls for all the *DROSHA* perturbation experiments.

**Fig.1.**
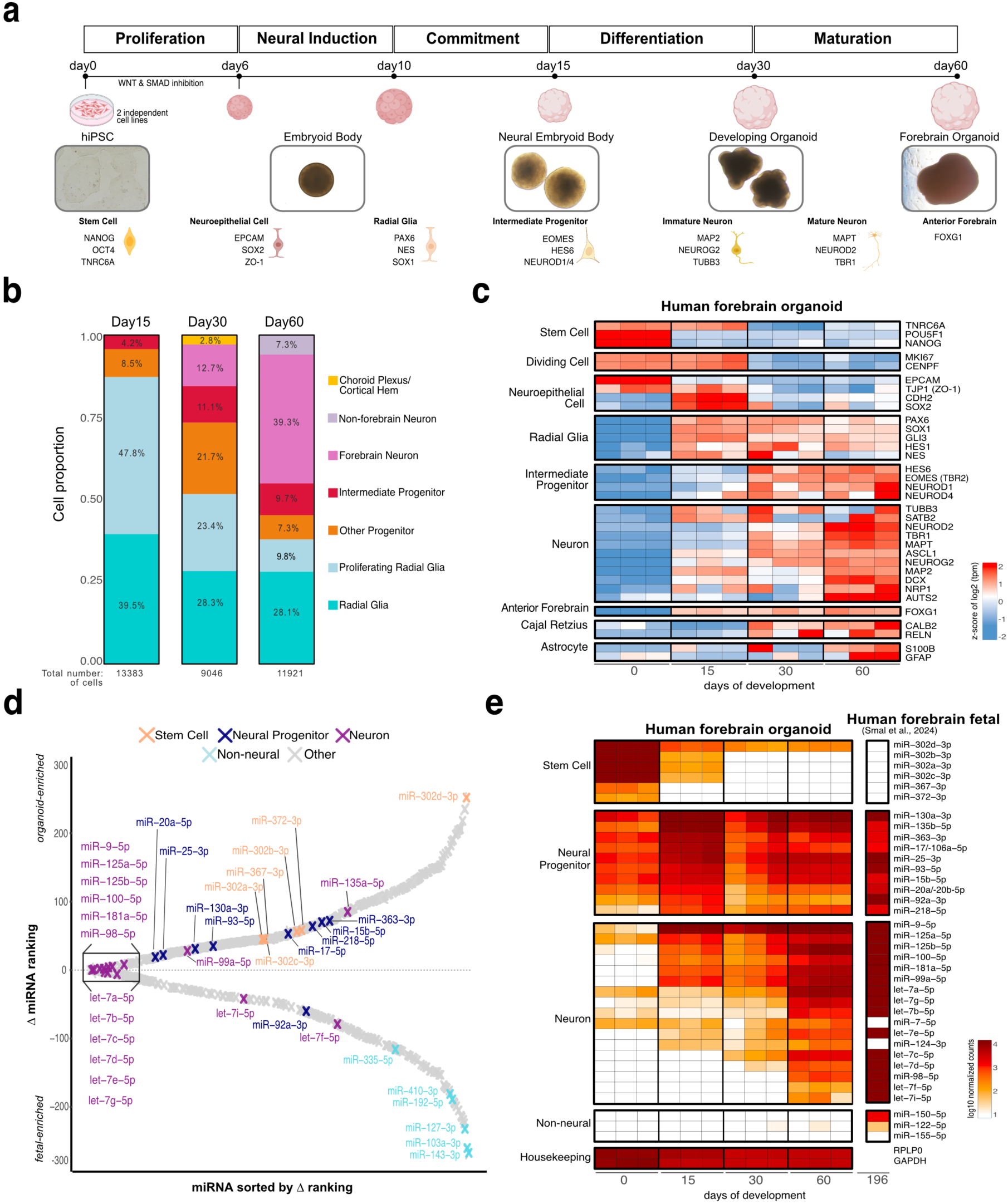
Neurodevelopmental mRNA and miRNA signatures are present in human brain organoids. **a.** Forebrain organoids recapitulate the sequential cell type transitions of early human brain development. The forebrain organoid generation protocol is initiated from hiPSC (=human induced pluripotent stem cell; “Methods”), across two independent cell lines. Brightfield images at different organoid developmental stages are shown, together with an overview of key cell transitions and corresponding marker genes. **b.** The cellular composition of organoids shifts from neural stem cells toward neurons, over time. Proportions of cell types across organoid development, from single cell RNA-sequencing analysis. Cluster annotation at each time point is shown. Day15=pool of 5 organoids; day30 and day60=pool of 3 organoids. Data derived from hiPS cell line1. **c.** Marker genes of major brain cell types are activated and silenced in the expected temporal order across organoid development. Gene expression quantified by bulk RNA sequencing, with expression values displayed as z-scores of log2 tpm (transcripts per million; “Methods”). Each column represents one biological replicate (Day15=pool of 6 organoids, day30 and day60= pool of 3 organoids). n = 3 for hiPS cell line1. **d.** The neuronal miRNA profile of 60-day organoids closely resembles that of the human fetal forebrain. Comparison of miRNA expression rankings in 60-day old organoids (nCounter) and in human fetal forebrain tissue (small RNA-seq from Smal et al., 2024). miRNA ranking difference (organoid rank − fetal rank), with positive and negative values respectively indicating organoid-enriched and fetal-enriched miRNAs. The dotted line at zero marks equal ranking (highest similarity) between the two systems. Crosses represent a single miRNA and are color-coded according to miRNA category. Selected miRNAs are labeled as representative examples of each category. **e.** miRNA signatures reflect the progression of human neurodevelopment. Expression of selected miRNAs split by cell type in human brain organoids and fetal brain. Direct quantification using nCounter (“Methods”), with expression values displayed as log10 normalized counts of miRNAs. Left heatmap: expression of miRNAs across organoid development (“Methods”), where each column represents one biological replicate (Day15=pool of 6 organoids, day30 and day60= pool of 3 organoids). n = 3 for hiPS cell line1. Right heatmap: expression of miRNAs in human fetal forebrain (195 days post-conception) (Smal et al., 2024).

**Fig.2.**
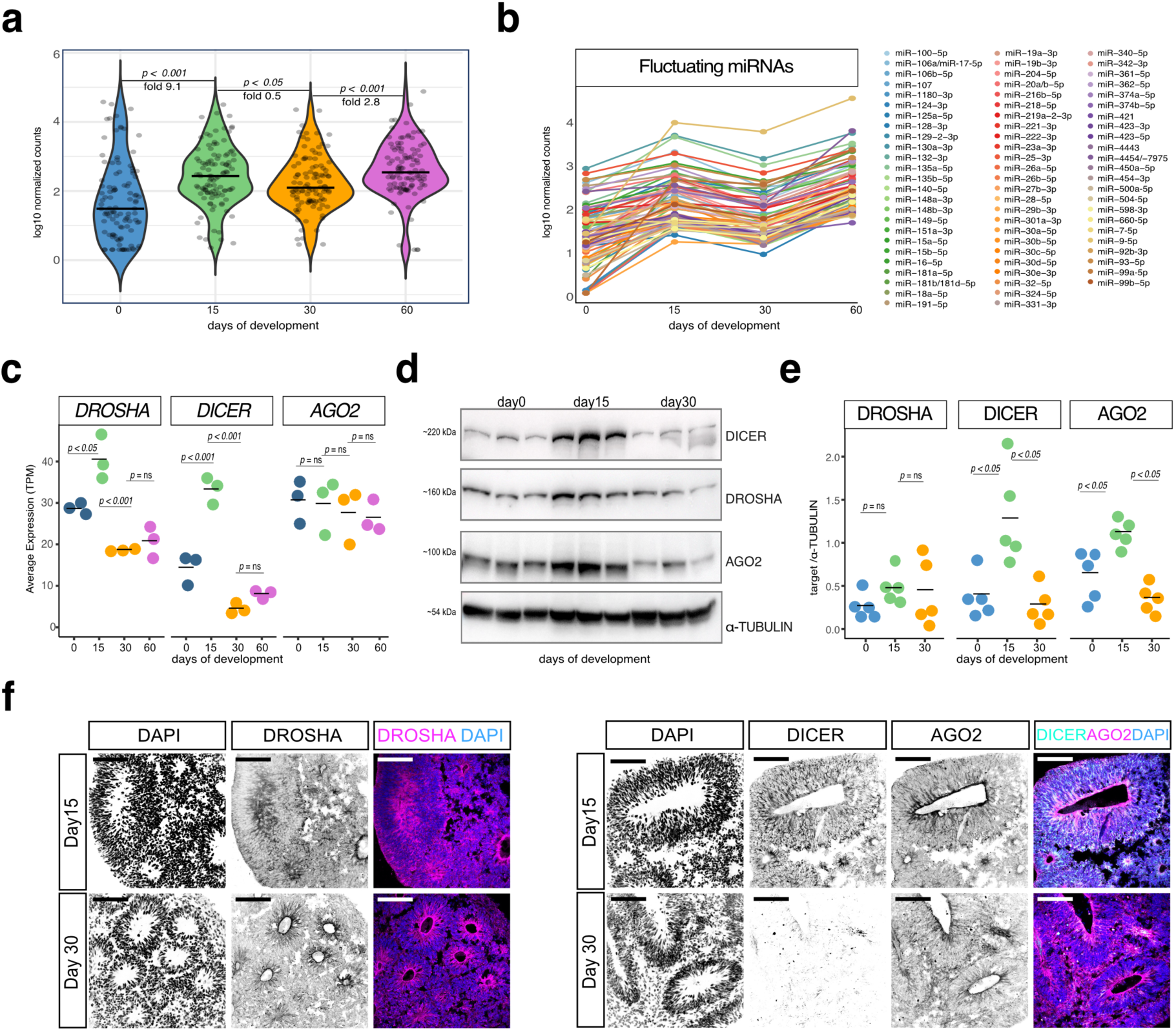
Expression dynamics of miRNAs and biogenesis machinery highlights specific developmental stages. **a.** Global miRNA expression changes dynamically over organoid development. Direct quantification using nCounter (“Methods”). Each dot represents the average log10 normalized counts (filtering: ≥ 100 counts at least at one stage) deriving from three biological replicates. (Day15=pool of 6 organoids, day30 and day60= pool of 3 organoids). n = 3 for hiPS cell line1. Median line and fold are displayed. Significance assessed using Wilcoxon rank-sum test. **b.** A subset of miRNAs fluctuates over development, suggesting stage-specific regulatory roles. Direct quantification using nCounter (“Methods”). Each line represents the average log10 normalized counts of three biological replicates (Day15=pool of 6 organoids, day30 and day60= pool of 3 organoids). n = 3 for hiPS cell line1. **c.** Transcripts of miRNA biogenesis machinery- DROSHA, DICER and AGO2- change dynamically across organoid development. Quantification from bulk RNA-sequencing (Day15=pool of 6 organoids, day30 and day60= pool of 3 organoids, “Methods”). Each dot represents an individual biological replicate. n = 3 for hiPS cell line1. Horizontal bars represent the mean. Holm-Bonferroni adjusted p-values are shown. **d.** DROSHA, DICER and AGO2 protein levels peak at commitment (day15). Western blot at different time points of organoid development, with α-TUBULIN serving as loading control. Each lane represents an individual biological replicate (Day15=pool of 8 organoids, day30 = pool of 3 organoids, “Methods”). n = 3 for hiPS cell line1. **e.** Quantification by densitometry of DROSHA, DICER and AGO2 proteins confirms their peak at commitment. Each dot represents an individual biological replicate (n=5 for hiPS cell line1, from two independent batches). Each protein is normalized to α-TUBULIN. Horizontal bars represent the mean. Holm-Bonferroni adjusted p-values are shown. **f.** DROSHA, DICER and AGO2 proteins are ubiquitously distributed throughout the organoid tissue. Immunofluorescence images (n=3, for hiPS cell line1) of 15 and 30 day-old organoids. Nuclei are stained by DAPI. Scale bar= 100 µm.

### 4. *DROSHA* perturbation experiments

*DROSHA* perturbation was first validated in hiPSCs and early EBs (**Fig. 3a-c; Suppl. Fig. 5a-c; Suppl. Fig. 6a-d**). Given the relatively long half-life and stability of miRNAs (Gantier et al., 2011; Guo et al., 2015), doxycycline (3 µg/mL) was administered for 3 days prior to downstream analyses. HiPSCs were seeded into 12-well plates in E8 Flex medium supplemented with ROCK inhibitor. Doxycycline treatment was initiated the same day during the first medium change (**Suppl. Fig. 5a**). In the EB differentiation protocol, dox was added from day 3 to day 6 (**Suppl. Fig. 6a**). To evaluate the perturbation penetrance, flow cytometry experiments were performed on EBs. Three independent pools, each composed of 80 doxycycline-treated single EBs, and three separate pools, each composed of 20 untreated control EBs, were collected. Pooled EBs were next washed with PBS, dissociated into single cells using TryplE and resuspended in E6 media. Dissociated cells were then centrifuged, resuspended in PBS and filtered through a 35 µm nylon mesh strainer cap fitter on flow cytometry tubes. Samples were kept on ice until sorting analysis. Flow cytometry was carried out using an LSR Fortessa™ X-20 cell analyser with HTS (BD, Biosciences). Data acquisition and analysis were performed using BD FACSDiva software (v.8.02). GFP-positive cells were sorted, collected, and stored at -80°C until RNA extraction (**Suppl. Fig. 6b-d**). For the perturbation experiments conducted in organoids, samples were treated with dox (3 µg/mL) at each media change until the day of collection, with main downstream analyses conducted at day 30 (**Fig. 3d**). In the neural induction (“early”) and neural differentiation (“late”) perturbation conditions, dox treatment was initiated either prior to commitment (day 5) or during commitment (day 9). GFP expression was routinely monitored by fluorescence microscopy. Untreated control organoids were generated in parallel using identical conditions, except dox administration.

**Fig.3.**
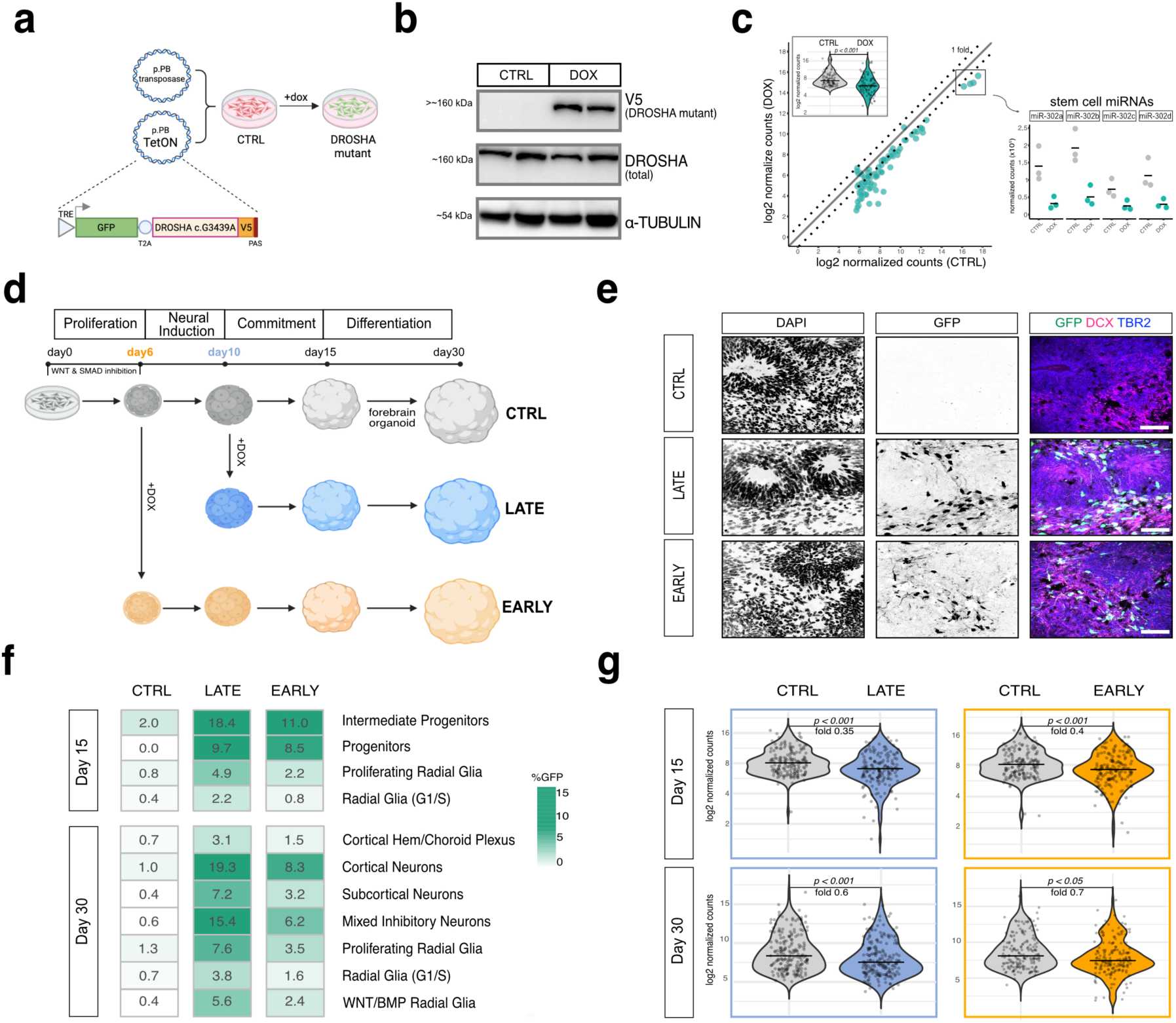
Expression dynamics of miRNAs and biogenesis machinery highlights specific developmental stages. **a.** A doxycycline-inducible dominant-negative DROSHA system enables temporal control of miRNA biogenesis. A TetOn construct, encoding TRE (tetracycline responsive element), GFP and DROSHA mutant (c.G3439A tagged with V5) was stably integrated in the hiPSCs, through PiggyBac (PB) transposase. Upon doxycycline (DOX) addition, the mutant protein is co-expressed with GFP via a T2A self-cleaving peptide, allowing live tracking of perturbed cells. **b.** DROSHA mutant protein is expressed upon DOX treatment. Western blot for total DROSHA and V5 (DROSHA mutant) proteins in control (CTRL) and perturbed (DOX) hiPS cells (“Methods”). α-TUBULIN serves as loading control. Each lane is a biological replicate for hiPS cell line 1. **c.** DROSHA perturbation (DOX) broadly reduces miRNA levels in hiPSCs, with the stem cell-specific miR-302 family among the most depleted (zoomed-in). Direct quantification using nCounter (“Methods”). Each dot represents the average log2 normalized counts (≥ 50) for three biological replicates of hiPS cell line 1. Dotted lines represent a log2 fold change of ±1. Insert: global shift in miRNA distribution, with median line displayed. Significance assessed using Wilcoxon rank-sum test. **d.** Experimental design of DROSHA perturbation time windows in human forebrain organoids. Organoids were DOX-treated from day 6 (“EARLY” = pre-neural commitment) and from day 10 (“LATE” = during neural commitment). Most downstream analyses were conducted on 30-day old samples. **e.** DROSHA perturbation is mainly targeting neural progenitors and neurons in 30-day organoids. Immunofluorescence images of 30-day old control (CTRL) and perturbed (“EARLY” and “LATE”) organoids showing GFP (perturbed cells), TBR2 (intermediate progenitors) and DCX (early neurons). Scale bar= 50um. hiPS cell line1. **f.** DROSHA perturbation predominantly affects neural cells. Percentage of GFP positive cells within each cell type identified by single cell RNA-sequencing (“Methods”). GFP was quantified in 15 and 30-day old organoids control and perturbed (“EARLY” and “LATE”) conditions. **g.** miRNA depletion is sustained throughout organoid development. Distribution of miRNA expression levels (nCounter) at days 15 and 30 in control (CTRL) versus perturbed (LATE and EARLY). Each dot represents the mean log2 normalized count (≥50) from three biological replicates of hiPS cell line 1 (day15) and cell line 2 (day30). Fold change and significance (Wilcoxon rank-sum test) are indicated.

### 5. Immunofluorescence

Organoids were washed three times with PBS, fixed in 4% paraformaldehyde for 30–60 min (depending on their age and size) at 4 °C, and washed three more times with PBS. With an overnight incubation at 4 °C, organoids were let settle in 40% sucrose (in PBS) and then embedded in 13%/10% gelatin/sucrose. Tissue blocks were stored at -80 °C. For sectioning, blocks were processed into 10 µm thick sections, using a cryostat. Sections were placed on slides and stored at -80°C. To proceed with immunostaining, slides were incubated for 5 min with warm PBS to remove the embedding solution and additionally fixed with 4% PFA for 10 min. After three washes with PBS, sections were blocked and permeabilized in 0.1% Triton-X, 0.2% BSA, and 4% normal goat or donkey serum in PBS for 1 h. Organoid sections were subsequently incubated with primary antibodies overnight at 4°C in blocking solution. On the next day, they were washed three times for 10 min with a washing solution (PBS supplemented with 0.1% Triton-X and 0.2% BSA). Secondary antibody incubation was then performed at room temperature for 2 h, followed by three washes, each of 10 min, in washing solution. DAPI (4′,6-diamidino-2-phenylindole, final concentration 1 µg/ml) was added to the last wash, in order to stain the nuclei. Samples were mounted with Prolong Gold Antifade mounting media (Thermo Fisher, catalog no. P36930) and imaged with a Keyence BZ-X710 microscope and a Leica STELLARIS 8 confocal microscope. *Z*-stacks and final images processing was done using Fiji-ImageJ, version 1.53f51, Java 1.8.0_172 (64-bit).

### 6. Protein extraction and Western Blotting

Protein extraction was performed on both hiPSCs and organoids from different developmental stages. hiPSCs were rapidly washed with PBS and harvested using 70 μL of RIPA buffer (150 mM NaCl, 5 mM EDTA, 50 mM Tris, 1% NP-40, 0.5% sodium deoxycholate, and 0.1% SDS). Depending on their age and size, 3-8 organoids were pooled together, washed once with PBS, and collected in RIPA buffer. RIPA was previously supplemented with protease and phosphatase inhibitors (Sigma-Aldrich, catalog no. 04693132001). Organoid samples were further homogenized in silicate beads using a tissue homogenizer device (Precellys Evolution). HiPSC and organoid-containing tubes were then cooled on ice for 30 minutes and centrifuged at 14000 g for 20 minutes at 4 °C. Eventually, the supernatant, containing the extracted protein, was collected and quantified using the Pierce BCA assay (Thermo Fisher, catalog no. 23225). To proceed with Western Blotting (WB), 15-20 ug of protein was denatured at 70°C for 10 min and run on a 6% polyacrylamide gel at 150 V in Tris-glycine buffer, together with a protein ladder (Page Ruler Prestained Plus, Thermo Fisher catalog no. 26619). Subsequently, semidry blotting using the Bio-Rad Trans-Blot Turbo Transfer system onto PVDF membranes was performed (Trans-Blot Turbo Midi 0.2 µm PVDF Transfer Packs, Bio-Rad catalog no. 1704157). Membranes were next blocked in 5% skimmed milk TBS-T for 1 h and probed with primary antibodies (diluted into 5% milk TBS-T) overnight at 4 °C. Employed primary antibodies are listed in **Table 3**. The next day, membranes were washed three times (each wash done for 30 min) in TBS-T and incubated for 1 h with HRP-conjugated secondary antibodies. Membranes were then imaged using chemiluminescent detection reagents (Thermo Fisher, catalog no. RPN2235) on a Vilber Fusion FX7 Edge imager with the Evolution-CaptEdge Fusion FX Edge (v.18.09) software. Band intensities were quantified using ImageJ, version 1.53f51 (functions from Analyze/Gels toolbar).

### 7. RNA extraction

Depending on their age and size, 2-6 organoids were pooled together, washed once with phosphate-buffered saline (PBS), and collected into 300-500 µl of homemade TRIzol. Next, organoids were homogenized in silicate beads using a tissue homogenizer device (Precellys Evolution). RNA extraction was done using the Zymo Directzol RNA miniprep kit (Zymo, catalog no. R2051), including DNase I digestion.

### 8. Target gene expression quantification

#### 8.1. Quantitative real-time PCR (qPCR)

Depending on their age and size, 2-6 organoids were pooled together, washed once with phosphate-buffered saline (PBS), and collected into 300-500 µl of homemade TRIzol. Next, organoids were homogenized in silicate beads using a tissue homogenizer device (Precellys Evolution). RNA extraction was done using the Zymo Directzol RNA miniprep kit (Zymo, catalog no. R2051), including DNase I digestion. 200 ng of RNA were used for cDNA synthesis. For the reverse transcription (RT) reaction, Maxima H minus reverse transcriptase (Thermo Fisher, catalog no. EP0751) was used according to the manufacturer’s instructions with random hexamers as primers. For the qPCR reaction, 4 ng of diluted cDNA were combined with SYBR green master mix (Biozym, catalog no. 331416S) and 0.5 μM forward and reverse primers (**Table 1**). The qPCR plates were processed by Roche 96 Light Cycler under the following cycling conditions: 95 °C for 10 min, then 40 cycles of 95 °C for 15 s and 60 °C for 1 min with fluorescence reading, and a final melting curve step. qPCR results were imported and analyzed using the LightCycler Roche Software 96 SW 1.1. Target mRNA expression was normalized to *GAPDH,* and relative quantification was calculated using the comparative ΔΔCT method (Pfaffl, 2001).

**Table1.**
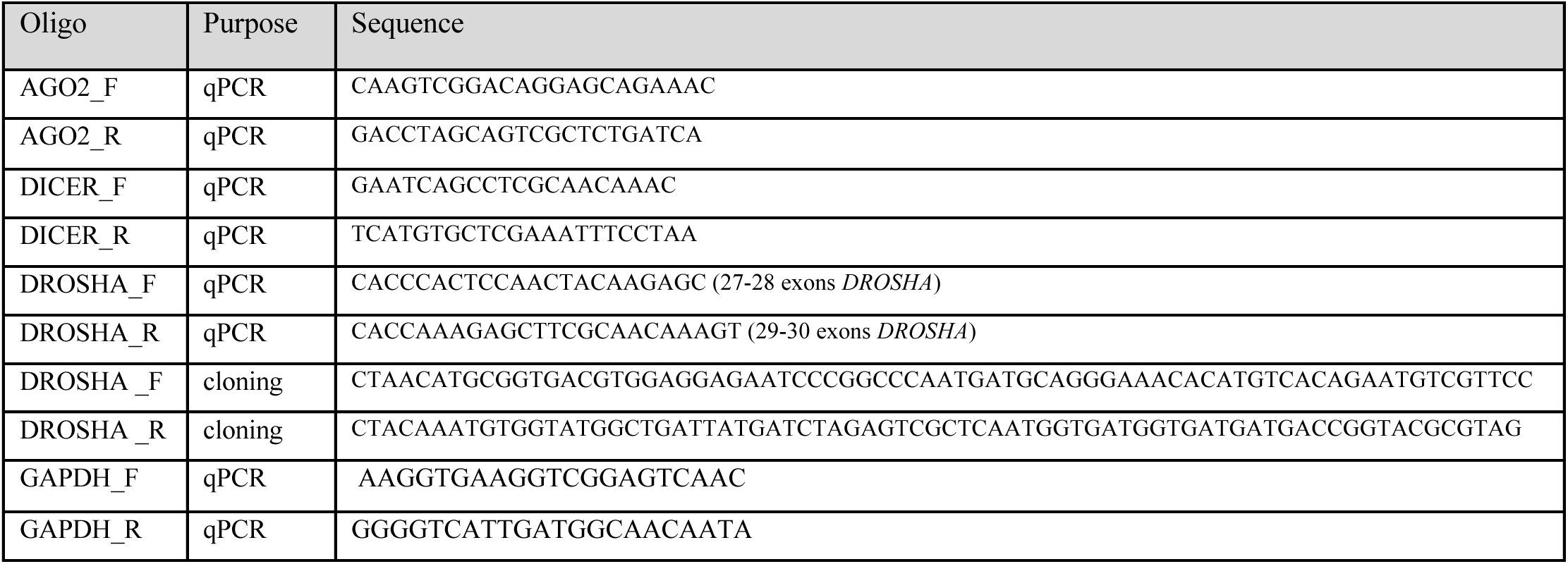
Primers.

#### 8.2. nCounter mRNA Expression Analysis

To quantify targeted single RNA molecules, we utilized the nCounter technology (Nanostring Technologies, USA), a probe-based, amplification-free method. Experiments were conducted according to the manufacturer’s protocol. In brief, 100 ng of pure total RNA was provided as input material and incubated with a 72-plex Core Tag set, a custom-made probe mix (reporter + capture probes), and the corresponding hybridization buffer. The reaction was carried out at 67°C for 18 h, followed by a cooldown step at 4°C for 10 minutes. The hybridization products were then diluted with RNase-free water to reach a final volume of 30 µl and injected into the nCounter gene expression cartridge (12-sample panel). Samples were processed using the nCounter SPRINT^TM^ Profiler instrument (Nanostring Technologies) for digital counting of RNA molecules. Background subtraction was next applied to raw counts of mRNA molecules and normalization done over the housekeeping controls *GAPDH, RPLP0*, and *TUBB*, using the nSolverTM Data analysis software Version 4.0 (Nanostring Technologies). For the analysis of mRNA expression across organoid development, raw data from all time points (day0 to day60) were normalized jointly.

### 9. miRNA expression quantification

#### 9.1. Taqman assays

To quantify individual miRNAs by qPCR, TaqMan assays were performed. For each reaction, 100 ng of total RNA was reverse transcribed in a 20 µL volume using the following components: SuperScript III (Invitrogen) (100 U/reaction), 20 U Ribolock (40 U/μl), 1 mM dNTPs, 1x first-strand synthesis buffer, 1x TaqMan RT primer per each miRNA of interest (**Table 2),** and nuclease-free water. RT was performed under the following cycling conditions: 30 min at 16°C, 30 min at 42°C, and 5 min at 85°C. Following cDNA synthesis, each sample was diluted 1:2 with nuclease-free water. qPCR was then conducted using TaqMan Universal Master Mix II, No UNG (Applied Biosystems, catalog no. 4440043). Target miRNA expression was normalized either to U6 or RNU48, and relative quantification was calculated using the comparative ΔΔCT method (Pfaffl, 2001).

**Table2.**
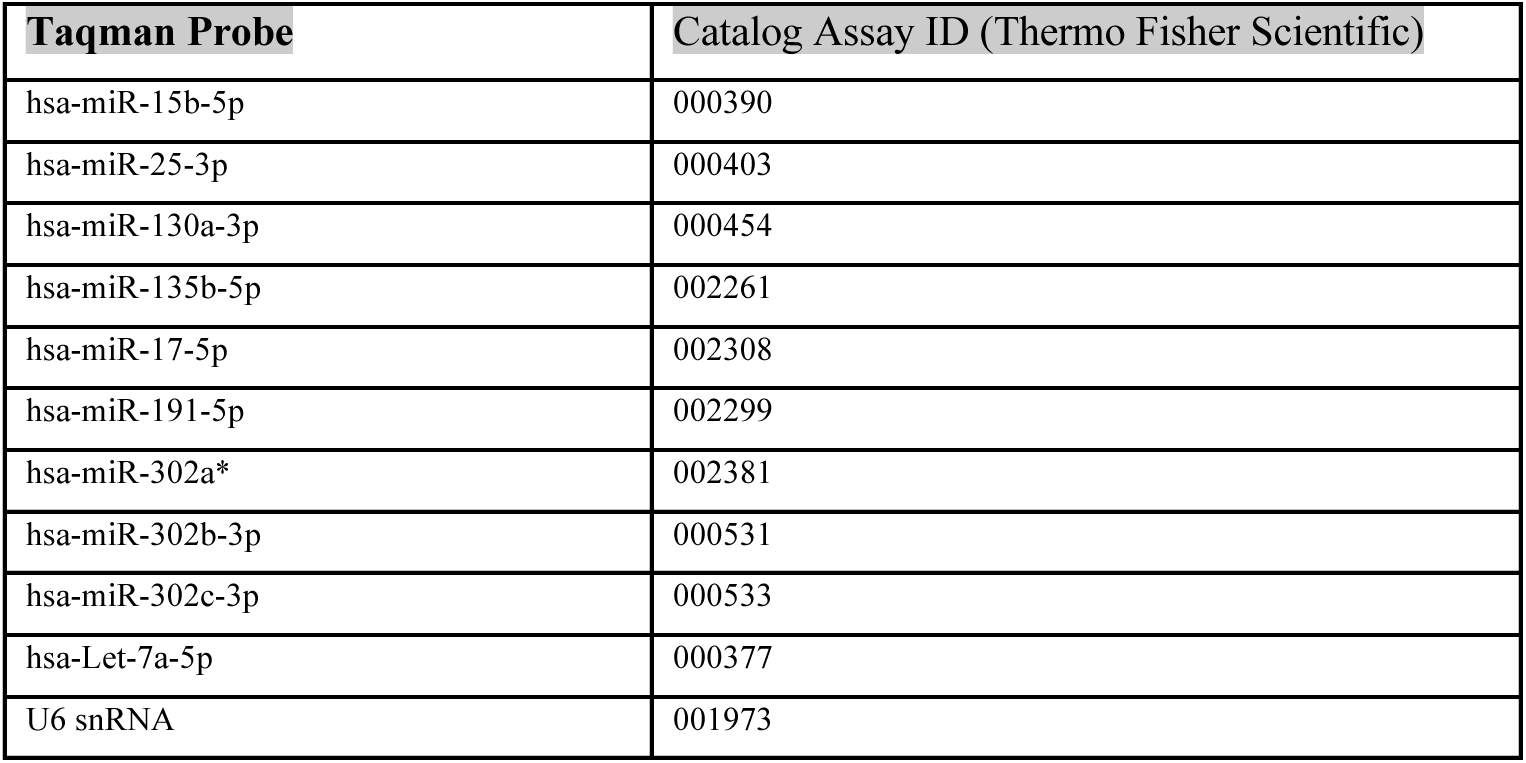
Taqman probes.

| Taqman Probe | Catalog Assay ID (Thermo Fisher Scientific) |
| --- | --- |
| hsa-miR-15b-5p | 000390 |
| hsa-miR-25-3p | 000403 |
| hsa-miR-130a-3p | 000454 |
| hsa-miR-135b-5p | 002261 |
| hsa-miR-17-5p | 002308 |
| hsa-miR-191-5p | 002299 |
| hsa-miR-302a* | 002381 |
| hsa-miR-302b-3p | 000531 |
| hsa-miR-302c-3p | 000533 |
| hsa-Let-7a-5p | 000377 |
| U6 snRNA | 001973 |

**Table3.**
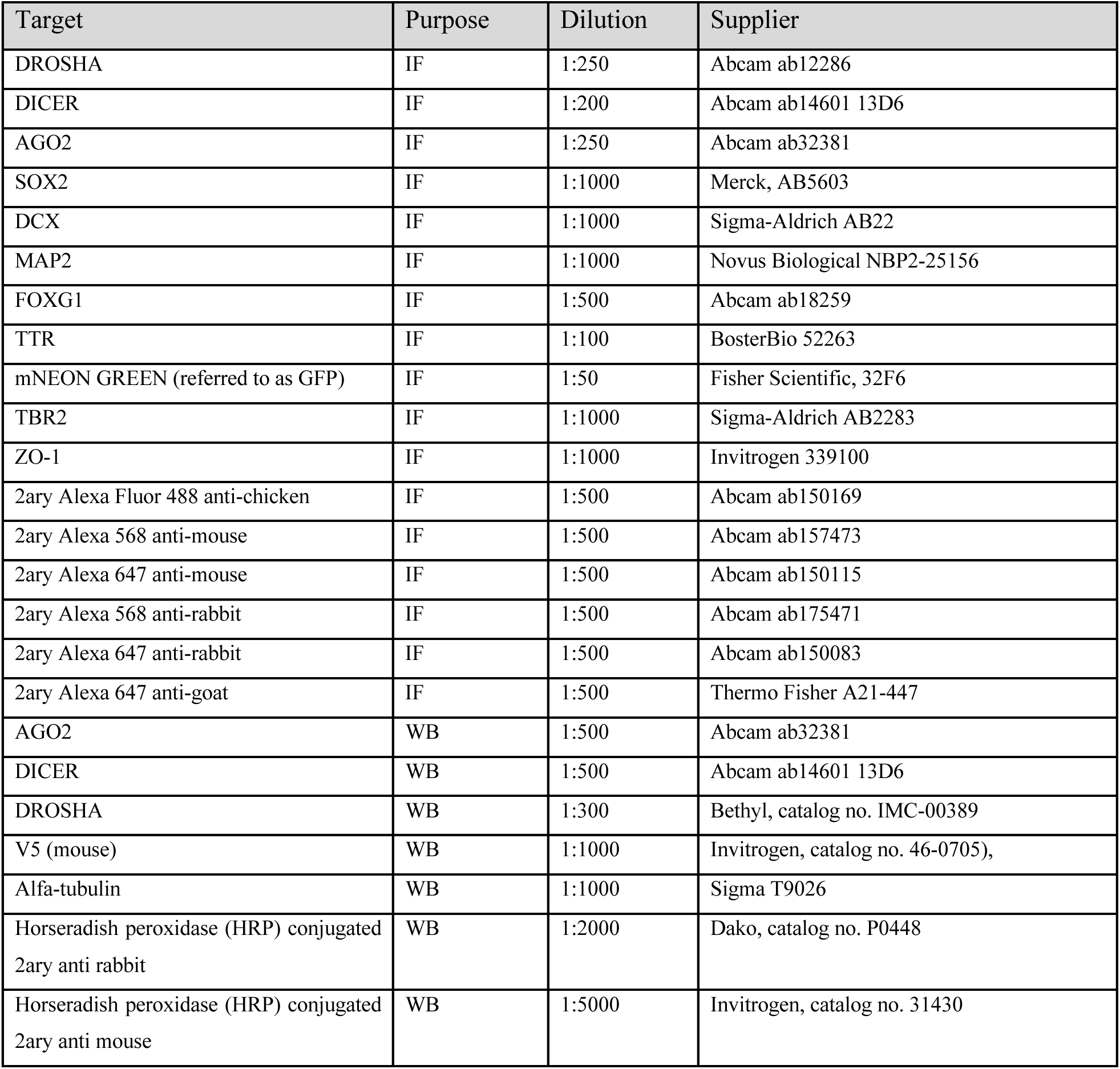
Antibodies.

| Target | Purpose | Dilution | Supplier |
| --- | --- | --- | --- |
| DROSHA | IF | 1:250 | Abcam ab12286 |
| DICER | IF | 1:200 | Abcam ab14601 13D6 |
| AGO2 | IF | 1:250 | Abcam ab32381 |
| SOX2 | IF | 1:1000 | Merck, AB5603 |
| DCX | IF | 1:1000 | Sigma-Aldrich AB22 |
| MAP2 | IF | 1:1000 | Novus Biological NBP2-25156 |
| FOXG1 | IF | 1:500 | Abcam ab18259 |
| TTR | IF | 1:100 | BosterBio 52263 |
| mNEON GREEN (referred to as GFP) | IF | 1:50 | Fisher Scientific, 32F6 |
| TBR2 | IF | 1:1000 | Sigma-Aldrich AB2283 |
| ZO-1 | IF | 1:1000 | Invitrogen 339100 |
| 2ary Alexa Fluor 488 anti-chicken | IF | 1:500 | Abcam ab150169 |
| 2ary Alexa 568 anti-mouse | IF | 1:500 | Abcam ab157473 |
| 2ary Alexa 647 anti-mouse | IF | 1:500 | Abcam ab150115 |
| 2ary Alexa 568 anti-rabbit | IF | 1:500 | Abcam ab175471 |
| 2ary Alexa 647 anti-rabbit | IF | 1:500 | Abcam ab150083 |
| 2ary Alexa 647 anti-goat | IF | 1:500 | Thermo Fisher A21-447 |
| AGO2 | WB | 1:500 | Abcam ab32381 |
| DICER | WB | 1:500 | Abcam ab14601 13D6 |
| DROSHA | WB | 1:300 | Bethyl, catalog no. IMC-00389 |
| V5 (mouse) | WB | 1:1000 | Invitrogen, catalog no. 46-0705), |
| Alfa-tubulin | WB | 1:1000 | Sigma T9026 |
| Horseradish peroxidase (HRP) conjugated<br>2ary anti rabbit | WB | 1:2000 | Dako, catalog no. P0448 |
| Horseradish peroxidase (HRP) conjugated<br>2ary anti mouse | WB | 1:5000 | Invitrogen, catalog no. 31430 |

#### 9.2 nCounter miRNA Expression Analysis

We utilized the nCounter miRNA Expression Assay (Human v3 miRNA CodeSet Kit 12 reactions, Nanostring Technologies, Seattle, USA), enabling digital counting of ∼800 human miRNAs. Experiments were conducted according to the manufacturer’s protocol. In brief, 100 ng of purified total RNA was annealed and ligated to unique oligonucleotide tags via target-specific bridge oligos. After a purification step, the tagged miRNAs were hybridized for 21 h at 65°C with customized reporter and capture probes. The following day, the hybridization products were directly injected into the nCounter gene expression cartridge (12-sample panel) and processed using the nCounterSPRINT™ Profiler instrument (Nanostring Technologies). Background subtraction was next applied to raw counts of miRNA molecules, and normalization was done over positive control probes of known concentration, employing the nSolver™ Data analysis software Version 4.0 (Nanostring Technologies). For the analysis of miRNA expression across organoid development, raw data from each time point (day 0 to day 60) was normalized jointly. In our samples, a substantial proportion of miRNAs included in the panel were either not expressed or expressed at very low levels, consistent with the context-specific nature of miRNA expression (**Suppl. Fig. 3a).** In **Fig. 1e** and **Suppl. Fig. 2b**, a subset of miRNAs was categorized into specific cell types (stem cell, neural progenitor, neuron, and non-neuronal) according to literature (Anokye-Danso et al., 2011; Nowakowski et al., 2018; Balzano et al., 2018; Jopling et al., 2005; Xiao and Rajewsky, 2009; O’Connell et al., 2007). In **Fig. 2a** and **Suppl. Fig. 4,** we analyzed the expression distribution of biologically relevant miRNAs (≥ 100 counts in at least one of the four developmental stages) for two independent hiPS cell lines. This filtered set comprised 106 miRNAs for hiPS cell line 1 and 186 miRNAs for hiPS cell line 2, corresponding respectively to ∼13% and ∼23% of the total probe panel. MiRNA expression in *DROSHA*-perturbed hiPSC and organoids was also quantified using the nCounter Assay (**Fig. 3c, g**, **Suppl. Fig. 5f** and **Suppl. Fig. 7a-b)**

### 10. Comparative analysis of miRNA expression in organoids and human fetal brain

miRNA expression profiles of forebrain organoids were compared with those of the human fetal brain (Smal et al., 2024). For the fetal samples, miRNA expression from the frontal, temporal, and parietal lobes was averaged and referred to as “anterior forebrain”, reflecting the regional identity toward which our organoids are patterned. The fetal sample corresponds to ∼ 28 postconception weeks (196 days), representing a more advanced developmental stage than our latest-stage organoids (day 60). Based on previous transcriptomic comparisons, 60-day-old brain organoids approximate ∼18 post-conceptional weeks of fetal development (Kelava and Lancaster, 2016) (**Suppl. Fig. 2c**). In Smal et al., raw small RNA-sequencing counts were normalized to total mapped reads, and miRNAs with ≥ 10 reads in at least one sample were retained, yielding 770 miRNAs. Of these, 475 miRNAs overlapped with the nCounter probe set used to quantify miRNAs in our organoids. miRNAs not in common were excluded from the analysis. miRNAs from both datasets were subsequently ranked by expression level, and rankings were compared between organoids and fetal anterior forebrain (**Fig. 1d-e**).

### 11. Bulk RNA sequencing experiments

#### 11.1. Library preparation

##### For control samples of hiPS cell line 1

Libraries were prepared from RNA of three biological replicates (*n* = 3) for each time point (days 0, 15, 30, and 60). For the organoid samples, each biological replicate consists of a pool of 6 organoids for day 15 and of 3 organoids for days 30 and 60. Samples were prepared as described in the “QuantSeq 3’mRNA-Seq V2 Library Prep Kit FWD with 12 nt Unique Dual Indices (UDIs)” protocol (Lexogen, catalog no. 191.96). In brief, 500 ng of RNA were reverse transcribed and adapter-ligated, followed by a pilot qPCR to determine the optimal PCR amplification. Library quality was assessed using Bioanalyzer DNA High Sensitivity and Qubit Fluorometer High Sensitivity (Invitrogen, catalog no. Q33226). Samples were sequenced on an Illumina NextSeq 500 using unique dual indexing with 1×84 bp.

##### For control samples of hiPS cell line 2 and *DROSHA*-perturbed samples of hiPS cell lines 1 and 2, before

library preparation, removal of ribosomal RNA was performed using the ribodepletion kit NEBNext® rRNA Depletion Kit v2 (human/mouse/rat kit) (New England Biolabs, catalog no. E7400X). Libraries were then prepared using the NEBNext Ultra II Directional RNA Library Prep Kit for Illumina (human/mouse/rat kit) (New England Biolabs, catalog no. E7760), according to the manufacturer’s instructions. In brief, 500 ng of RNA was fragmented and subjected to reverse transcription. The derived cDNA was ligated to sequencing adapters, followed by PCR amplification. Library quality and average size were evaluated using the Agilent Bioanalyzer DNA Chip. Libraries were quantified using Qubit. Eventually, libraries were pooled together and sequenced on an Illumina NovaSeq X Plus 10B lane, 200 cycles, in paired-end modality with 100 bp for read1 and read2.

#### 11.2. Data processing and analyses

Raw sequencing data (BCL files) were first converted to FASTA files and demultiplexed using bcl2fastq v2.20 for the identification of reads based on sample-specific barcodes. Quality control was performed using FastQC v. 0.12 (Andrews, 2010). Next, trimming of low-quality bases, poly-A tails, poly-G tails, and adapter sequences was carried out using Cutadapt v.5.0 (Martin, 2011), discarding low-quality and short (<30 bp) sequences. Proper sequence processing was verified with FastQC. The processed RNA-seq reads were mapped to the human genome assembly GRCh38.p13 (GCF_000001405.39) using STAR Aligner v.2.6.1a (Dobin et al., 2013). Default mapping parameters were used. Aligned reads were assigned to genes and quantified using annotations from GENCODE (release 41) and featureCounts v.1.6.4 (Liao et al., 2014). The resulting count table was used as input to conduct differential gene expression analysis (DEA). DEA analyses were performed in RStudio software with R v.4.4.1, using DESeq2 v.1.44.0 (Love et al., 2014), with default options and a unifactorial design (design = ∼ condition). Human gene annotation was obtained from the Ensembl database (GRCh38.p13, GCF_000001405.39). Data were filtered for significance using an adjusted p-value (FDR) threshold of < 0.05 and an absolute log₂ fold change cutoff > 1 (**Suppl. Fig. 1e**). For the analysis of gene expression changes across organoid development (from day 0 to day 60), principal component analysis (PCA) was first performed to assess the overall quality of the dataset (**Suppl. Fig. 1d**). Sequencing counts were normalized using DESeq2 v.1.44.0 within RStudio with R v.4.4.1, and global expression changes were visualized on volcano plots with filtering by baseMean>100 (**Suppl. Fig. 1e** and **Suppl. Fig. 7c)** or on MA plots (**Fig. 4a)** with filtering by baseMean>10. Expression changes of genes of interest were visualized on heatmaps (**Fig. 1c**, **Fig. 4f** and **Suppl. Fig. 7d):** sequencing counts were first normalized on gene length and converted to log tpm (transcripts per million), which were next z-scored. For **Fig. 1c**, marker gene annotation was based on Hendricks et al. (2024). For **Fig. 4f**, WNT and BMP gene signatures’ categorization into ligands, receptors, agonists, antagonists, and effectors was based on Klaus and Birchmeier (2008) and Clevers and Nusse (2012) for WNT and on Lim et al. (2000) and Wang et al. (2014) for BMP. Transcriptomic changes between control and *DROSHA*-perturbed organoids underwent a gene set enrichment analysis (GSEA). This was focused on GO (Gene Ontology) BP (Biological Process) using the enrichGO function in clusterProfiler (v4.0.5) (Yu et al., 2012). Multiple testing correction was applied using the Benjamini-Hochberg (BH) method, with a p-value cutoff of 0.05 to identify significantly enriched GO terms (**Fig. 4b, d**).

**Fig.4.**
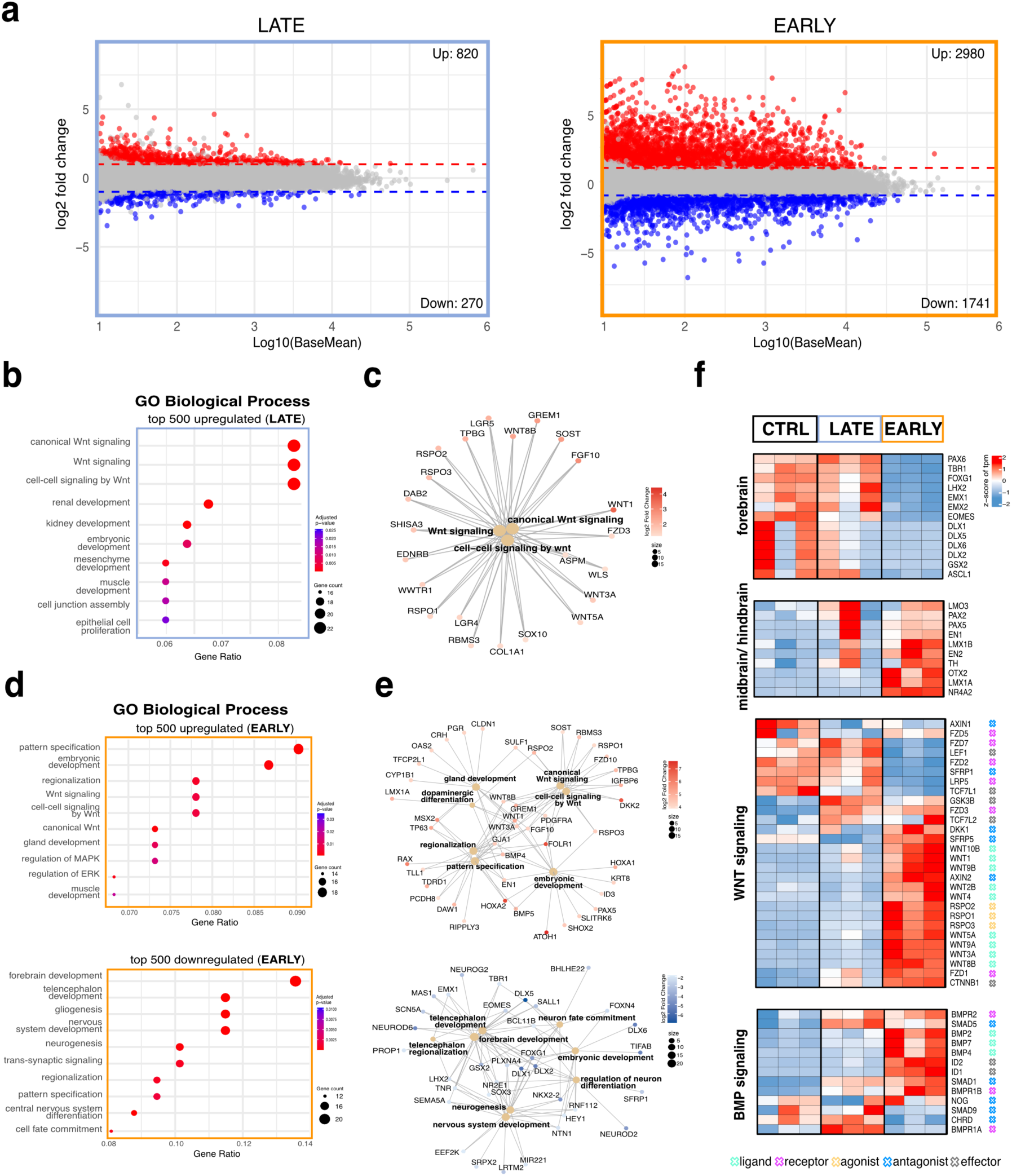
MiRNAs regulate forebrain patterning within a critical developmental window. **a.** EARLY perturbation causes a stronger transcriptional response than LATE perturbation. MA plot shows differentially expressed genes in: “LATE” (left) and “EARLY” (right) relative to control (CTRL) at 30-day organoids (“Methods”). Upregulated genes are shown in red, downregulated in blue; grey dots are not significant (adjusted p-value > 0.05, |log2FC| > 1). Grey: not significant. Data from three biological replicates per condition (*n* = 3; each pool consists of 3 organoids), from hiPS cell line 1. **b.** WNT signaling is the dominant pathway upregulated in LATE miRNA perturbation. Dot plot shows the top 10 enriched Gene Ontology (GO) Biological Process (BP) terms among the 500 most upregulated genes in “LATE” versus control (CTRL) organoids. Dot size corresponds to the number of associated genes. Color gradient represents the enrichment significance. P-values: Benjamini-Hochberg-corrected GSEA test. **c.** Primary upregulated genes driving WNT pathway enrichment in LATE organoids. Network plot connects enriched GO terms for “LATE” versus control (CTRL) organoids (beige nodes) with upregulated genes (outer nodes). The color intensity and the size of each gene node correspond to the magnitude of upregulation. **d.** Early perturbation activates both WNT signaling and broad developmental patterning programs, while suppressing forebrain-specific gene expression. Dot plot shows the top 10 enriched Gene Ontology (GO) Biological Process (BP) terms among the 500 most upregulated genes in “EARLY” versus control (CTRL) organoids. Dot size corresponds to the number of associated genes. Color gradient represents the enrichment significance. P-values: Benjamini-Hochberg-corrected GSEA test. **e.** Upregulated genes in EARLY-perturbed organoids converge on WNT-driven patterning, while downregulated genes reflect loss of forebrain and neurogenic identity. Network plot connects enriched GO terms for “EARLY” versus control (CTRL) organoids (beige nodes) with upregulated (top) and downregulated (bottom) genes (outer nodes). The color intensity and the size of each gene node correspond to fold-change magnitude. **f.** Forebrain transcription factors are progressively silenced, while WNT/BMP signaling components are induced in EARLY-perturbed organoids. Heatmap shows gene signatures of forebrain identity, midbrain/hindbrain identity, WNT and BMP signaling in control (CTRL), “LATE” and “EARLY” 30-day organoids. Expression values are displayed as z-scores of log tpm (transcripts per million; “Methods”). For WNT and BMP signatures, gene symbols are annotated by functional role (ligand, receptor, agonist, antagonits, effector). Each column represents one biological replicate (*n* = 3; each pool consits of 3 organoids) from hiPS cell line1.

### 12. 5’ RNA sequencing

#### 12.1. Library preparation

Three libraries of 30-day-old organoids of “control,” “early,” and “late” conditions (hiPS cell line 1) were prepared. For each library, three organoids were pooled together and dissociated to single-cell preparations using a papain- and trypsin-based method from the Neural Tissue Dissociation Kit (P), according to the manufacturer’s instructions (Miltenyi Biotec, catalog no. 130-092-628). In brief, collected organoids were first chopped using a surgical blade and incubated consecutively with two enzymatic mixes. After neutralization of the enzymatic reactions with a stop solution (DMEM + FBS), filtering through a 70 μm cell strainer, and centrifugation (300 g, 4°C, 10 min) were performed. The cell pellet was next resuspended in PBS + 0.04% BSA and filtered through a 40 μm cell strainer. The number and viability of single dissociated cells were then assessed using automated cell counting with trypan blue. Single cells were next fixed with the Fixation of Cells & Nuclei for Chromium fixed RNA profiling kit (10x Genomics, catalog no. PN-1000414), according to the manufacturer’s instructions. Fixed cells with viability > 95% were directly used for library preparation, which was kindly carried out by the Genomics Facility at BIMSB/MDC, using the 10x 5’ GEM Single Cell RNA-seq kit (10x Genomics, catalog no. PN-1000265). Cell suspensions were adjusted to 1,600 cells/μl as input, aiming at a cell recovery of ∼ ∼20,000 cells. Libraries were prepared following manufacturer’s instructions and sequenced on Illumina NovaSeq X Plus 10B lane in paired-end modality with 100 bp for read1 and read2, 100 cycles.

#### 12.2. 5’ RNA sequencing data processing

Raw sequencing data (BCL files) were first converted to FASTA files and demultiplexed using bcl2fastq v2.20, for the identification of reads based on sample-specific barcodes. Quality control was performed using FastQC v.0.12 (Andrews, 2010). Next, trimming of low-quality bases, poly-A tails, poly-G tails, and adapter sequences was carried out using Cutadapt v.5.0 (Martin, 2011), discarding low-quality and short (<30 bp) sequences. Proper sequence processing was verified with FastQC. The processed RNA-seq reads were mapped to the human genome assembly GRCh38.p13 (GCF_000001405.39) using STAR Aligner v.2.6.1a (Dobin et al., 2013). Default mapping parameters were used. The resulting BAM alignment files were converted to normalized bigWig tracks using ‘bamCoveragè from the ‘deepTools’ package v.3.5.6 (Ramírez et al., 2016) and visualized using the Integrative Genomics Viewer Version 2.4.3 (Robinson et al., 2011).

### 13. Single-cell RNA-seq cell

#### 13.1. Single-cell RNA-seq data generation

Single cell RNA sequencing was conducted on organoids from hiPS cell line 1. For 15-day-old samples, 6 organoids were pooled together and then dissociated to single cells using TrypleE and filtered through a FACS tube. For 30- and 60-day old samples, 3 organoids were pooled and the dissociation was conducted using 30a papain and trypsin-based method from the Neural Tissue Dissociation Kit (P), according to manufacturer’s instructions (Miltenyi Biotec, catalog no. 130-092-628). In brief, collected organoids were first chopped using a surgical blade and incubated consecutively with two enzymatic mixes. After neutralization of the enzymatic reactions with a stop solution (DMEM and % FBS), filtering through a 70 μm cell strainer and centrifugation (300 g, 4°C, 10 min) was performed. The cell pellet was next resuspended in PBS containing 0.04% BSA and filtered through a 40 μm cell strainer. Number and viability of single dissociated cells were next assessed using automated cell counting with trypan blue. Single cells were subsequently fixed with Fixation of Cells & Nuclei for Chromium-fixed RNA profiling (10x Genomics, PN-1000414), according to the manufacturer’s instructions. Fixed cells were stored at -80 °C until processing. Single cell preparations with viability > 90% and ∼ 1×10^6 cells were used for library preparation and sequencing with the probe-based “Chromium Human Transcriptome Probe Set v1.0.1” according to manufacturer’s instructions (10x Genomics). To the 53,957 probes included in the panel, we added customized probes targeting neon green (for simplicity, “GFP”) (**Table 4).** Libraries were sequenced using the Seq388 / NovaSeq X Plus 10B lane, 100 cycles from Illumina.

**Table4.**
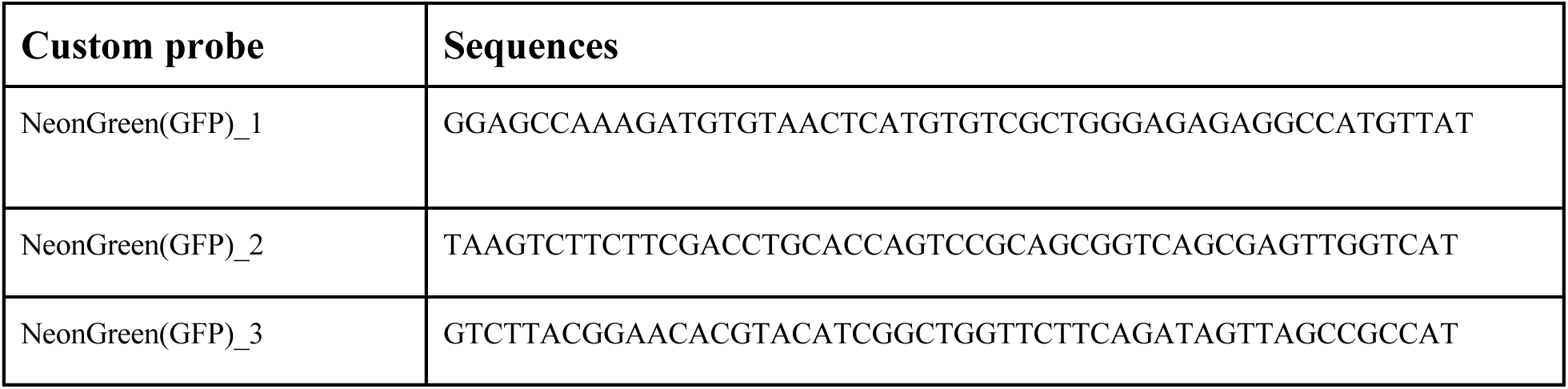
Custom probes for 10x single cell RNA sequencing.

#### 13.2. Single-cell RNA-seq data processing

FASTQ files were generated using Cellranger v.8.0.1 (‘Cellranger mkfastq’) and aligned to the GRCh38 human genome. A reference genome (’cellranger count’) to generate digital gene expression matrices with default parameters. After quality filtering, we collected 67474 single-cell transcriptomes across 6 samples from hiPSC cell line 1 (two biological replicates of each condition). Then, we performed low-dimensional embedding and clustering of control and *DROSHA*-perturbed samples. Filtered gene expression matrices were imported in R v4.4.3 for downstream processing using Seurat v5.3.0. Single time points (days 15, 30, and 60) were merged into independent Seurat objects, and cells with fewer than 100 detected genes and more than 10% mitochondrial transcripts were removed. Doublets were identified using scDblFinder, and cells with a doublet score higher than 0.5 were removed. Raw gene expression counts were normalized.

#### 13.3. Single-cell RNA-seq cell type identification

The top 2000 variable genes were used to compute 50 principal components and UMAP coordinates. Clusters were identified using FindNeighbours and FindClusters (resolution = 0.2, dims=1:10). Cluster annotation was performed by examining top 50 marker genes per cluster obtained from FindMarkers, supported by known marker gene expression and label transfer scores from Rybak-Wolf et al., 2023. Label transfer was performed using the standard Seurat v5.3.0 pipeline (FindTransferAnchors normalization.method= ‘SCT’, dims=1:10, non.method=’rann’), eps=0.5), as previously shown in Rybak-Wolf et al., 2023.

#### 13.4. Differential gene expression

To identify differentially regulated genes between “EARLY” and “LATE,” we performed pseudobulk analysis. For each annotated cluster, pseudobulk expression profiles were generated by summing raw gene expression counts across all cells belonging to the same biological sample within each cluster. Deprecated gene annotations were excluded from downstream analysis. For pairwise condition comparison, differential gene expression analysis was performed using the DESeq2 package (v1.38.3), and for each comparison, a DESeq2 dataset was created from the pseudobulk count matrix. MA plots were generated using ggplot2. Genes with low expression values (baseMean < 10) were filtered out, and genes with log₂FC > 1 or -1 were highlighted.

#### 13.5. Cell proportions quantifications

For each sample, the proportion of cells belonging to each cluster was calculated as the percentage of total cells, based on cluster assignment, using the standard Seurat v5 pipeline. These proportions were summarized and visualized with bar plots as a percentage of change from the control condition.

#### 13.6 Regional identity scoring

To assess the regional identity of cells across conditions, gene module scores were computed using the AddModuleScore function from Seurat v5.3.0. Two gene sets were defined based on established regional markers: a forebrain program (LHX2, EMX2, EMX1, FOXG1, EOMES, TBR1, PAX6, GSX2, ASCL1) and a midbrain/hindbrain program (OTX2, LMO3, LMX1A, EN1, EN2, TH, NR4A2, PAX2, PAX5, LMX1B). Scores were computed using 24 expression bins and 100 control genes randomly sampled from each bin (nbin = 24, ctrl = 100). Scores were projected onto UMAP coordinates separately for each condition (Control, Early, Late). Negative scores are render in grey to highlight cells with above-background program activity.

#### 13.7. Regional marker gene expression across conditions

To compare the expression of genes from the forebrain and midbrain/hindbrain programs across conditions at single-gene resolution, the percentage of cells expressing each marker gene (raw count ≥ 1) was computed per condition using SCT-normalized counts. Pairwise scatter plots were generated comparing the percent-positive values between conditions (Control vs Early, Control vs Late, Early vs Late).

#### 13.8. Morphogen quantification

The average expression of morphogen gene sets was quantified for Cortical Hem/Choroid Plexus cluster across condition using SCT-normalized counts. For each gene set, average expression was calculated per gene and per cluster and condition. Expression values were then row-wise z-scored to facilitate cross-gene comparison of relative expression patterns.

#### 13.7. CellChat

Intercellular communication analysis was performed using CellChat package v2 to identify and visualize morphogen signaling pathways, specifically *WNT, BMP, FGF*, and *NOTCH*. Data were first split by experimental condition: “CONTROL,” “EARLY,” and “LATE,” and a separate CellChat object was constructed and analyzed for each condition using cluster and condition groupings as cell identities. Interaction probabilities between cell populations were inferred using CellChat’s built-in signaling pathway databases and statistical framework. Interaction networks for selected pathways were visualized using circle plots.

### 14. Candidate miRNA identification and target predictions

To identify candidate miRNAs, the following filtering criteria were applied to our nCounter datasets of miRNA expression in control and early-perturbed organoids: 1) miRNAs expressed at >100 counts, in at least one developmental time point, in the control condition were retained; 2) miRNAs downregulated in 30-day-old organoids, upon “early” perturbation (log₂FC ≥ 0.5), were kept. Moreover, those two parameters had to be met consistently across the two independent cell lines. Applying this filtering yielded 6 candidate miRNAs of interest: miR-15b, miR-93, miR-221, miR-362, let-7b, and miR-149-5p. To select their target transcripts, predictions from TargetScan (https://www.targetscan.org/vert_80/ McGeary et al., 2019) and miRDB (https://mirdb.org/, Liu and Wang, 2019; Chen and Wang, 2020) were integrated, considering unique targets predicted by both tools. To those, additional stringency was imposed based on TargetScan scoring metrics, focusing on transcripts with conserved 8-mer or 7-mer-m8 seed matches and strong predicted repression. Gene expression profiles of the top 200 predicted targets for each of the candidate miRNAs were examined in our bulk RNA-seq datasets (baseMean > 50). Expression changes of targets versus non-target genes were visualized using cumulative distribution function (CDF) plots. Differential gene expression (log₂ fold changes of “early” vs. control) between predicted targets and background control was tested for significance using a two-sided Kolmogorov-Smirnov test. A shift toward global target upregulation was observed for 5 (miR-15b, miR-93, miR-221, miR-362, and let-7b) of the 6 miRNA candidates.

### 15. miRNA gain-of-function experiments

miRNA gain-of-function experiments were conducted in a 2D culture system of neural progenitor cells (NPCs). HiPSCs were seeded on 2% Geltrex-coated 6-well plates and cultured for 4 days in neural forebrain induction media (E6 supplemented with 5 µM XAV, 10 µM SB and 0.1 µM LDN). Subsequently, 250,000 cells were seeded per well onto Geltrex-coated 12-well plates and transfected on days 4 and 9. For transfection, Lipofectamine RNAiMAX (Thermo Fisher, catalog no. 13778150) was diluted in Opti-MEM according to the manufacturer’s instructions. The transfection mix contained either 50nM scramble control constructs- miRIDIAN microRNA Mimic Negative Control #1 (CN-001000-01-05, Dharmacon) and #2 (CN-002000-01-05, Dharmacon) for “CTRL” and “EARLY” conditions, or 50nM of a pooled combination of the five candidate miRNA mimics: miRIDIAN miRNA mimic hsa-miR-15b-5p (C-300587-05-0002, Dharmacon), hsa-miR-93-5p (C-300512-07-0002, Dharmacon), hsa-miR-221-3p (C-300578-05-0002, Dharmacon), hsa-miR-362-5p (C-300664-05-0002, Dharmacon ), and hsa-let-7b-5p (C-300476-05-0002, Dharmacon). At day 4, medium was exchanged 7h after transfection with neural induction media without XAV supplementation, with doxycycline (3 µg/mL) added where needed. At day 9, medium was replaced 7h after transfection with neural differentiation medium (COM1). Thereafter, the medium was changed every other day. From day 16 onward, cells were maintained in neural maturation media (COM2). Cells were collected for downstream analyses at days 10, 15 and 30.

## Tables-Methods

The following primers were purchased from IDT (PrimeTime™ qPCR Primers): *AXIN2, BMP2, EMX2, FOXG1, FZD3, RSPO2, WNT1, WNT4, WNT3A*

## Results

### 1. Brain organoids recapitulate mRNA programs and conserved neuronal miRNA signatures of early human brain development

To model human brain development *in vitro*, we generated stem cell-derived forebrain organoids by combined inhibition of WNT and SMAD signaling, followed by an optimized protocol based on Walsh et al. (2024) (**Fig. 1a**).

Between days 6 and 10, neural induction occurs as defined by the emergence of neural progenitor cells (radial glia), which express the key marker genes *SOX2, PAX6,* and *NES*. From day 10 to day 15, cells start to commit toward cortical neuronal lineages, giving rise to intermediate progenitors and immature neurons, with neuronal maturation reached by day 60 (**Fig. 1a**). At day 15, organoids form ZO1+ ventricle-like structures, which are surrounded by SOX2+ radial glial cells. As differentiation progresses, TBR2+ intermediate progenitors and DCX+ and MAP2+ immature neurons emerge and localize radially to the progenitor structures (**Suppl. Fig. 1a, c**).

To systematically characterize the cellular composition of the organoids over time, we performed single-cell RNA sequencing (Methods, **Fig. 1b** and **Suppl. Fig. 1b-c)**. At day 15, radial glia cells were the dominant cell type, representing ∼85% of the total cell population. By day 30, progenitor populations expanded and immature neurons appeared, together comprising approximately 40% of the total cells. By day 60, neurons became the predominant population, accompanied by a marked reduction in radial glia cells (**Fig. 1b** and **Suppl. Fig. 1b**).

Based on these characteristics, we focused on four key developmental time points: day 0 (pluripotent stem cell stage), day 15 (neural commitment), day 30 (onset of neurogenesis), and day 60 (neuronal maturation). For a more robust molecular characterization, we conducted bulk RNA sequencing (Methods, **Fig. 1c** and **Suppl. Fig. 1d-e**). Principal Component Analysis (PCA) revealed a clear separation between hiPSCs and brain organoids, with the latter aligning sequentially along a developmental trajectory (**Suppl. Fig. 1d**). This was consistent with the differential gene expression observed across time points (**Suppl. Fig. 1e**).

We next examined the temporal expression of key developmental markers (**Fig. 1c** and **Suppl. Fig. 2a**). Pluripotency and proliferation genes were highly expressed at the earliest time points and decreased rapidly upon neural induction (**Fig. 1c** and **Suppl. Fig. 2a).** By day 15, neural progenitor markers confirmed proper acquisition of a neuroectodermal fate (**Fig. 1c** and **Suppl. Fig. 2a**), followed by increased expression of intermediate progenitor markers, *TBR2* (EOMES) and *HES6,* and immature neuronal markers, *DCX* and *MAP2,* as development advanced (**Fig. 1c**). Day 30 shows a highly plastic stage in cell fate specification, marked by ongoing neurogenesis and still relatively high levels of progenitor proliferation. By day 60, distinct neuronal subtypes became detectable, including immature deep-layer projection neurons, expressing *NEUROD4, NEUROD1,* and *MAPT;* and callosal projection neurons, marked by *SATB2* expression (**Fig. 1c**). Additionally, markers for Cajal-Retzius cells (*RELN* and *CALB2*) and astrocytes *(GFAP* and *S100B)* were detected at this stage. High *FOXG1* expression by day 60 confirmed organoid patterning towards an anterior-dorsal fate (**Fig. 1c** and **Suppl. Fig. 2a**).

In summary, these complementary approaches demonstrate that our forebrain organoids recapitulate expected gene expression programs and cellular diversity characteristic of early human neurogenesis, providing a foundation for subsequent miRNA functional studies (**Figs. 1a-c**).

We next profiled miRNA expression, utilizing a high-throughput, probe-based Nanostring platform (Methods) that enables direct quantification of ∼800 microRNA species. First, we compared miRNAs expressed in 60-day-old forebrain organoids with human fetal anterior forebrain (post-conceptional day 196) (Smal et al., 2024) (**Fig. 1d-e**). We assessed global similarities in miRNA profiles by ranking from high to low all overlapping miRNAs expressed in both systems (Methods) (**Fig. 1d)**. Notably, neuron-specific miRNAs showed the highest similarities between organoid and fetal samples, highlighting conserved regulatory programs (**Fig. 1d**, purple). Specifically, members of the let-7 family, miR-9, miR-125, miR-100, miR-181a, and miR-98 displayed the same expression pattern in both systems (**Fig. 1d**, purple). On the other hand, miRNAs known to regulate the differentiation from stem cells to neural progenitors (Anokye-Danso et al., 2011; Gioia et al., 2014; Jauhari et al., 2018; Yang et al., 2021) ranked higher in organoids (**Fig. 1d**, orange and dark blue), highlighting different developmental stages- more advanced in the fetal tissue compared to the organoids. Conversely, a group of miRNAs, “non-neural”, is found expressed uniquely in the fetal samples (**Fig. 1d**, light blue). This group comprises miRNAs associated with immune activation, such as miR-103-3p (Chen et al., 2020; Huang et al., 2021), and endothelial cell differentiation, such as miR-143-3p (Wang et al., 2020; González-López et al., 2024), consistent with the absence of microglia and endothelial cells in our model.

We next focused on a subset of miRNAs, associated with specific cell types (stem cell, neural progenitor, neuron, and non-neuronal) according to literature (Anokye-Danso et al., 2011; Jönsson et al., 2015; Nowakowski et al., 2018; and Balzano et al., 2018) (**Fig. 1e** and **Suppl. Fig. 2b).** We observed dynamic shifts in miRNA expression mirroring the developmental progression of human brain organoids, with the latest stage (day 60) most closely resembling miRNA profiles from human fetal forebrains (**Fig. 1e**).

As expected, in organoids, the expression of stem cell-specific miRNAs such as the miR-302/367 cluster (Anokye-Danso et al., 2011) declined after induction, consistent with loss of pluripotency, whereas (also as expected) neural-specific miRNAs, such as miR-130a, miR-135b, miR-17, and miR-9 (Nowakowski et al., 2018), increased over time (**Fig. 1e**). By day 60, we observed upregulation of all let-7 family members, along with miR-125, miR-181a, miR-124, and miR-7, consistent with the acquisition of neuronal identity. Negative controls, i.e. non-neuronal miRNAs, including the liver-specific miR-122 (Jopling et al., 2005) and immune cell-specific miRNAs (Xiao and Rajewsky, 2009) such as miR-150 (Monticelli et al., 2005) and miR-155 (Vigorito et al., 2013), exhibited very low or undetectable expression (**Fig. 1e** and **Suppl. Fig. 2b**).

Altogether, our data show that miRNAs can serve as robust molecular markers of organoid differentiation, with neuronal miRNA expression patterns in maturing organoids closely matching those observed in developing human fetal forebrain.

### 2. Developmental stage specific expression of miRNAs and their biogenesis machinery

We quantified global miRNA expression (Methods) during organoid development (**Fig. 2a** and **Suppl. Fig. 4a**). In both independent cell lines, we found that most miRNAs show a fluctuating expression pattern over time, suggesting stage-specific regulatory roles (**Fig. 2a** and **Suppl. Fig. 4a**). In particular, we observed peaks at days 15 and 60, with day 15 representing the neural commitment phase and day 60 reflecting the stage of neuronal maturation. This group of miRNAs comprises a mixture of neural progenitor, neuron-associated, and uncharacterized miRNAs (**Fig. 2b**). Other smaller groups follow different expression dynamics: 1) peak at day 0 (stem cell-enriched families) 2) peak at day 60 (neuron-enriched) (**Suppl. Fig. 3c**).

To relate these patterns to miRNA processing, we examined the expression of core components of the miRNA biogenesis machinery. *DROSHA* and *DICER* were highly transcribed at day 15, coinciding with the peak in miRNA expression and neural commitment (**Fig. 2c, Suppl. Fig. 3d** and **Suppl. Fig. 4b).** In contrast, *AGO2* transcripts remained stable across all time points (**Fig. 2c** and **Suppl. Fig. 4b**). At the protein level, all three miRNA biogenesis machinery components were strongly expressed at day 15 (**Fig. 2d-f** and **Suppl. Fig. 3f-g**). At days 30 and 60, miRNA machinery protein levels did not change, indicating an early transient boost at day 15 followed by stabilization (**Suppl. Figs. 3f-g**). In an independent cell line, this early peak of the machinery at day 15 extended over time, until day 30 (**Suppl. Fig. 4c-d**). Immunostaining at day 15 and day 30 revealed a ubiquitous distribution of DICER, DROSHA and AGO2 proteins within and around the neuroepithelial loops (**Fig. 2f**), with pronounced downregulation of DICER at day 30 in both neural progenitors and neurons (**Fig. 2f**).

Single-cell transcriptomics analysis from day 15 to day 60 confirmed that the biogenesis transcripts are broadly expressed across all time points and cell types, reflecting shifts in organoid composition: high at day 15 in radial glia and enriched in neurons at day 60 (**Suppl. Fig. 3e**).

In summary, we found that the abundance of miRNAs and their biogenesis machinery fluctuate during organoid differentiation, with a peak at the stage of neural commitment. These results suggest that miRNA production is regulated within a critical temporal window that coincides with neural specification (**Fig. 2**).

### 3. Overexpression of an inducible, dominant-negative *DROSHA* dampens mature miRNA expression

To test the hypothesis of a temporal requirement for miRNA function during early forebrain organoid development, we engineered two independent hiPSC lines to express a dominant-negative *DROSHA* mutant (c.G3439A, p.E114K) (Rakheja et al., 2014; Torrezan et al., 2014), along with a GFP reporter, upon doxycycline (dox) administration (**Fig. 3a**). Employing this inducible mutant, which specifically blocks maturation of miRNAs, allowed us to overcome the lethality phenotype of a constitutive knockout, as described in previous studies (Dicer: Bernstein et al., 2003; Ago2: Morita et al., 2007; DGCR8: Wang et al., 2007, reviewed in detail by Alberti and Cochella, 2017).

Administering dox for 3 days resulted in an average 4-fold upregulation of *DROSHA* mRNAs (**Suppl. Fig. 5a-b)** and the specific expression of the mutant protein (**Fig. 3b** and **Suppl. Fig. 5c**). Interestingly, expression of the DROSHA mutant altered the levels of other components of the miRNA biogenesis pathway, particularly an increase of DICER levels (**Suppl. Fig. 5d-e**). Consistently with previous reports, disruption of a core miRNA biogenesis factor can trigger compensatory upregulation of others within the pathway (Kim et al., 2016).

As expected for the dominant-negative *DROSHA* mutant, we observed a significant global downregulation of miRNAs upon perturbation across both hiPSC lines (average of **∼**2.6-fold reduction) (**Fig. 3c** and **Suppl. Fig. 5f-g)**. Stem cell-enriched miRNAs, particularly the miR-302 family, were among the most significantly reduced (**Fig. 3c**, right**; Suppl. Fig. 5f,** right and **Suppl. Fig. 5g)**.

We also examined whether the perturbation affected non-canonical, transcription-related functions of *DROSHA* as proposed by Gromak et al. (2013) (**Suppl. Fig. 5h**). Using 5’ RNA sequencing (Methods), we detected no changes between control and perturbed organoids in the transcription start sites usage for previously validated *DROSHA*-regulated genes, suggesting that the transcriptional activity at these loci was unaffected by the mutant (**Suppl. Fig. 5h**).

To assess perturbation penetrance, we sorted GFP⁺ cells via flow cytometry in early embryoid bodies (EBs). Approximately 30% of cells were GFP⁺, indicative of a mosaic effect (**Suppl. Fig. 6a-c, e**). Enriching for GFP⁺ cells significantly augmented miRNA downregulation, resulting in an average 4-fold reduction (**Suppl. Fig. 6d**). Considering the known miRNA long half-life (Gantier et al., 2011; Guo et al., 2015) and the observed peak of miRNA production at day 15 (**Fig. 2**), we next perturbed miRNA biogenesis at two distinct developmental stages: 1) during neural commitment, starting at day 10 (“late”) (**Fig. 3d**, blue) and 2) prior to neural commitment, starting at day 6 (“early”) (**Fig. 3d**, orange). Of note, following *DROSHA* perturbation, we did not observe any morphological abnormalities: organoids retained normal size and shape and developed neuroepithelial loop structures comparable to controls (**Fig. 3e** and **Suppl. Fig. 6e**). Interestingly, transgene-expressing GFP+ cells localized predominantly outside neuroepithelial loops and co-localized with TBR2+ intermediate progenitors and DCX+ maturing neurons (**Fig. 3e** and **Suppl. Fig. 6e**). This suggested that *DROSHA* perturbation predominantly affected neurogenic lineages, likely due to the hUBC promoter driving the TetON system, known to be more active in neurons (Wilhelm et al., 2011).

Single-cell RNA-seq further supported this observation: at day 15, neuronal progenitors showed stronger GFP expression (19% - 28% of the total number of cells in the “early” and “late” conditions, respectively) compared to radial glia (3% - 7% of the total number of cells in the “early” and “late” conditions, respectively) (**Fig. 3f**). Total GFP expression was higher in “late” (35% at day 15 and 62% at day 30) organoids compared to “early” (23% at day 15 and 27% at day 30) (**Fig. 3f)**. Correspondingly, DROSHA mutant protein levels were more elevated in “late” organoids than “early” ones (**Suppl. Fig. 6f**).

In summary, we established an inducible system to perturb miRNA biogenesis in forebrain organoids, allowing the temporal dissection of miRNA function during human development.

### 4. MicroRNAs are critical for cell fate acquisition and tissue patterning during early neurogenesis

Next, we quantified miRNA expression in *DROSHA*-perturbed organoids derived from two independent hiPSC lines (**Fig. 3g** and **Suppl. Fig. 7a-b**).

We observed consistent and significant global miRNA dysregulation in both “early” and “late”, at both days 15 and 30 (average log2 fold change = -1.4 at day 15 and average log2 fold change = - 0.7 at day 30) (**Fig. 3g**). However, while “late” showed a uniformly downregulated miRNA population at day 30 (**Suppl. Fig. 7a-b, blue**), the dysregulation in “early” was driven by a specific subset of miRNAs (**Suppl. Fig. 7a-b, orange**), suggesting distinct underlying molecular effects. We focused subsequent analyses on 30-day-old organoids, as this stage captures most cell types and key lineage transitions (**Fig. 1b-c** and **Suppl. Fig. 2a**).

Total bulk transcriptomic profiling revealed a global divergence between “early” and “late” for both cell lines (**Fig. 4a** and **Suppl. Fig. 7c)**. In “late”, only a limited number of genes were significantly altered (**Fig. 4a**, left and **Suppl. Fig. 7c,** left), while “early” organoids showed widespread transcriptional dysregulation (**Fig. 4a**, right and **Suppl. Fig. 7c,** right). Subsequent gene ontology analysis revealed a robust enrichment of the WNT signaling pathway in both conditions (**Fig. 4b-e**, top 500 upregulated). Uniquely in the “early” perturbation, we found: 1) enrichment of terms associated with regionalization, pattern specification, and dopaminergic differentiation (**Fig. 4d-e**, top); and 2) downregulated pathways related to forebrain development, neuronal fate commitment, telencephalon development, and neurogenesis (**Fig. 4d-e**, bottom).

We further examined key transcriptional signatures belonging to these biological processes. Forebrain-specific transcription factors, including *FOXG1, EMX1,* and *EMX2*, were extensively suppressed only in “early”, indicating disrupted regional identity, despite the forebrain-directed differentiation (**Fig. 4f** and **Suppl. Fig. 7c,** right). In contrast, midbrain/hindbrain markers such as *NR4A2, TH,* and *EN2* were strongly induced in “early” (**Fig. 4f** and **Suppl. Fig. 7c,** right). We further observed strong WNT pathway activation in “early”, characterized by the consistent upregulation of WNT ligands and agonists, together with a mixed regulation of WNT receptors and effectors (**Fig. 4f** and **Suppl. Fig. 7c,** right).

Given the well-established crosstalk between WNT and BMP signaling during early neurogenesis (Kasai et al., 2005; Kleber et al., 2005), we examined the expression of key BMP pathway genes. We observed consistent upregulation of BMP ligands and agonists, with a mixed regulation of other pathway components (**Fig. 4f)**. By contrast, FGF and NOTCH signaling were not remarkably altered in either “early” or “late” (**Suppl. Fig. 7d**).

Together, these findings suggest that only pre-commitment perturbation (“early”) led to large-scale reorganization of key patterning programs. This divergence aligns with transcriptional upregulation of WNT and BMP pathway components and downstream targets.

To further investigate these hypotheses, we performed single-cell RNA sequencing (Methods). We identified seven major cell populations, which we annotated based on cell-type-specific marker genes (Methods) (**Fig. 5a** and **Suppl. Fig. 8a-b**). These included radial glia, cortical excitatory and inhibitory neurons, cortical hem/choroid plexus cells, and subcortical neurons (**Fig. 5a** and **Suppl. Fig. 8a-b)**. The global cell-type distribution reflected the expected developmental trajectory (**Suppl. Fig. 8b**).

**Fig.5.**
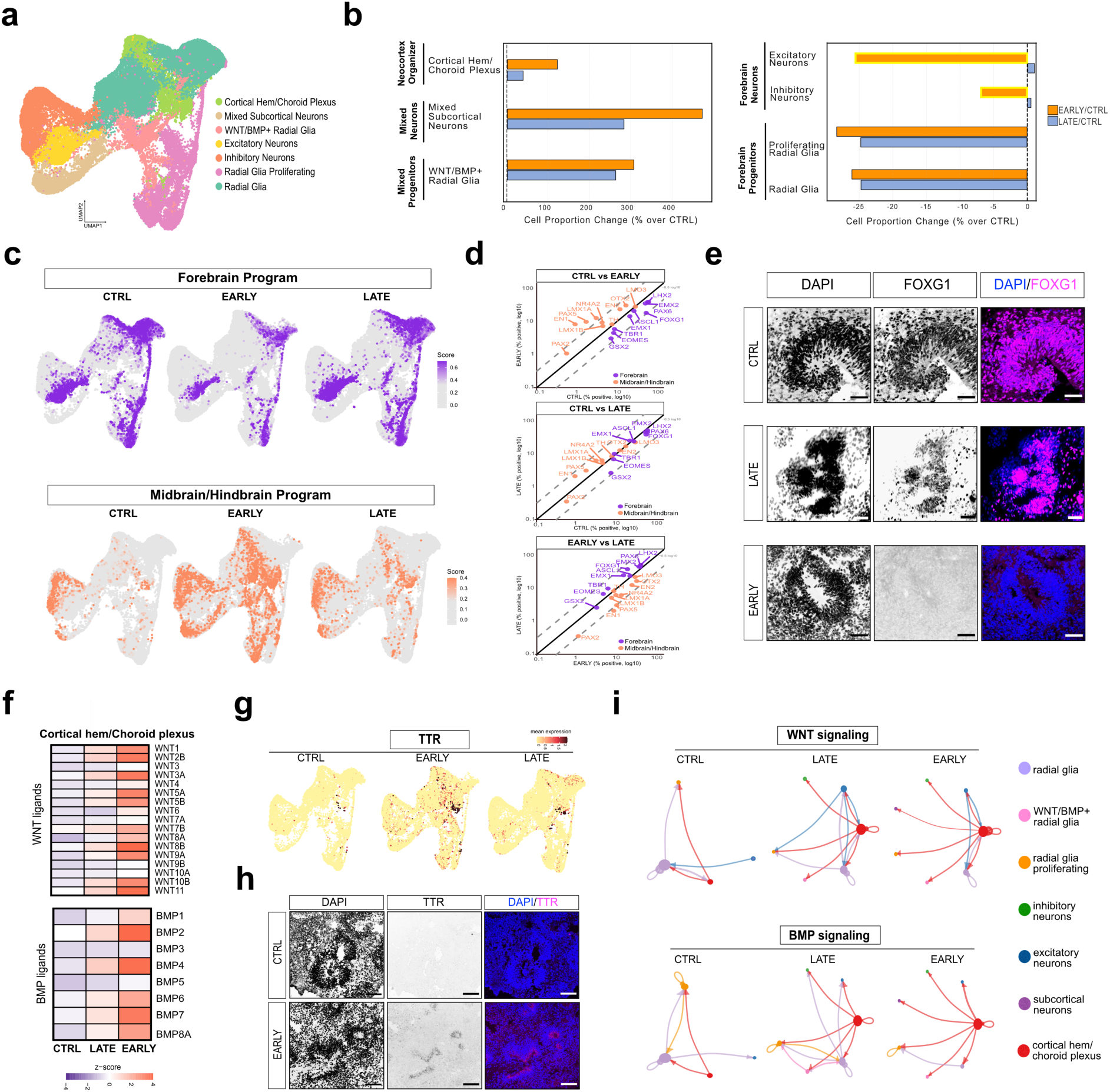
MiRNA disruption causes WNT/BMP imbalance redirecting organoid patterning and neural identity. **a.** Single-cell transcriptomic landscape of 30-day organoids across all three conditions (CTRL, “EARLY”, “LATE”). Uniform manifold approximation and projection (UMAP) visualization of single cell RNA-sequencing data with annotated clusters, revealing the major neural progenitor and neuronal populations present (“Methods”). Cluster annotation is shown. Two independent biological replicates per condition from hiPSC line1. **b.** Early and late miRNA perturbation shift cortical neuronal composition in opposite directions and forebrain progenitor populations are differentially affected between the two perturbation windows. Bar plots show the percentage of cell type proportion changes for “EARLY” (orange) and “LATE” (blue) relative to control (CTRL); “Methods”). **c.** Forebrain identity is progressively lost while midbrain/hindbrain identity expands with “EARLY” miRNA perturbation. UMAPs show per-cell activity scores of Forebrain (purple) or Midbrain/Hindbrain (orange) gene programs across conditions (CTRL, “EARLY”, “LATE”; “Methods”). Grey cells score at/or below background. Data from two independent biological replicates per condition. **d.** Individual regional marker genes shift expression in a program-consistent manner across conditions. Scatter plots compare the percentage of cells expressing each Forebrain (purple) and Midbrain/Hindbrain (orange) marker genes between conditions. Positive cells > 1 normalized count. Dashed lines represent ±0.5 log10 (∼3 folds). Data from two independent biological replicates per condition from hiPSC line1. **e.** FOXG1 protein, key forebrain transcription factor, is reduced in perturbed organoids. Immunofluorescence images of 60-day organoids stained for FOXG1 (magenta) and DAPI (blue; nuclei) across CTRL, “EARLY” and “LATE” conditions. Scale bar= 50um. **f.** WNT and BMP ligand expression is upregulated in the cortical hem/choroid plexus upon perturbation. Heatmap shows pseudobulk expression of WNT and BMP ligands (“Methods”), for cortical hem/choroid plexus cluster across conditions (CTRL, “EARLY”, “LATE”). Data from two independent biological replicates per condition. **g.** TTR, a choroid plexus marker, expands markedly in “EARLY” perturbed conditions. UMAP show per-cell TTR expression across conditions (CTRL, “EARLY”, “LATE”). Data from two independent biological replicates per condition. **h.** TTR protein confirms choroid plexus expansion in “EARLY” perturbed organoids. Immunofluorescence images of 60-day organoids stained for TTR (magenta) and DAPI (blue; nuclei). Scale bar = 100um. **i.** miRNA perturbation rewires WNT and BMP intercellular signaling networks. Cell communication analysis (CellChat; “Methods”) showing predicted signaling interactions among cell clusters for WNT and BMP pathways across all conditions (CTRL, “EARLY”, “LATE”). Node size reflects the number of signaling interactions per cluster. The thickness of the edges represents the strength of the communication between sender and receiver populations.

Interestingly, the overall composition changes were more pronounced in “early” compared to “late,” confirming the critical time dependency of the perturbation (**Fig. 5b** and **Suppl. Fig. 8c**). Specifically, we observed: 1) a substantial increase of cortical hem/choroid plexus cells (“early”: ∼120%; “late”: ∼30% over control); 2) an increase of subcortical neurons (“early”: ∼450%; “late”: ∼280% over control); and 3) a particular population of radial glial cells with high expression of *WNT* and *BMP* ligand genes (“early”: ∼300%; “late”: ∼270% over control) (**Fig. 5b** and **Suppl. Fig. 8c**).

While forebrain progenitors in “early” and “late” showed a similar decrease in cell proportion, their transcriptional signatures diverged extensively (**Fig. 5b** and **Suppl. Fig. 8d**). In these progenitors, “early” displayed upregulation of midbrain-hindbrain boundary markers — EN1, PAX5, DMBX1, FGF8, FGF17, and FOXC1. This suggests that the forebrain program is disrupted uniquely in “early”, consistent with posterior progenitor re-patterning (**Suppl. Fig. 8d**). This was further supported by the expansion of subcortical neurons (SLC17A7⁺/NR4A2⁺/CHAT⁺) at the expense of excitatory (TBR1⁺/vGLUT1⁺/RELN⁺) and inhibitory (GAD⁺/DLX⁺/SST⁺) cortical forebrain neurons (**Fig. 5b** and **Suppl. Fig. 8c**), suggesting a unique disruption of the forebrain program selectively in “early”.

Concordantly, subcortical neurons in “early” were enriched for posterior hindbrain (*HOXA2, CYP26C1, SPRY4)* and brainstem interneuron markers *(NPY, NOS1),* whereas “late” retained the dorsal forebrain identity (*TRB1, DLX2, SLC32A1 (vGAT), NEUROG1)* (**Suppl. Fig. 8d**). Importantly, cortical fate clusters showed strong depletion of forebrain progenitors in both “early” and “late” (“early”: ∼25-28%; “late”: ∼24% over control), but only “early” displayed the consequent reduction of cortical neurons (excitatory: ∼25%; inhibitory: ∼7%; **Fig. 5b** and **Suppl. Fig. 8c** and **f**). Together, this data revealed that the mispatterning occurring only in the “early” perturbation consists of the loss of cortical neurons and the increase of posterior markers (midbrain/hindbrain) (**Fig. 5b; Suppl. Fig. 8c and f**).

We next examined transcriptional activation of forebrain and midbrain/hindbrain programs. Uniquely in “early” we found an inhibition of the forebrain program (including *FOXG1, LHX2, PAX6, TBR1, EOMES, GSX2*, and *EMX1*) within the radial glial populations. with a simultaneous enrichment of midbrain/hindbrain program activity (including *EN1, EN2, LMX1B, LMX1A, NR4A2*, and *OTX2*) mostly across WNT/BMP+ radial glia and neuronal clusters **(Fig. 5c-d)**.

In agreement, we found that when the neuronal fate is fully defined (60-day-old organoids), the FOXG1 protein is visibly reduced uniquely in the “early” condition (**Fig. 5e**). Moreover, in “early” we detected a strong expansion of the cortical hem/choroid plexus (**Fig. 5b** and **Suppl. Fig. 8a**). The cortical hem is a known source of WNT and BMP morphogens (Furuta et al., 1997; Grove et al., 1998).

Thus, we investigated WNT and BMP ligand expression patterns across all cell populations (**Suppl. Fig. 8e)**. In “early” we observed a stronger activation of both *WNT* and *BMP* ligands localized in the cortical hem (**Fig. 5f** and **Suppl. Fig. 8e)**. Consistently with this, the choroid plexus marker *TTR* was substantially enriched at the RNA (**Fig. 5g**) and protein levels (**Fig. 5h**). We hypothesized that in “early” the high levels of WNT and BMP morphogens derived from the cortical hem organizer would imbalance the forebrain development, and consequently, this would lead to a shift in patterning. To investigate ligand-receptor interactions of WNT and BMP morphogens among cell types, we performed CellChat analysis (Methods). We observed that radial glia cells were the main senders and receivers of WNT and BMP communication in the control condition, whereas in the perturbed contexts (“early” and “late”), the main source of signaling shifted to the cortical hem and choroid plexus (**Fig. 5i**).

In summary, miRNA perturbation prior to neural commitment (“early”) induces WNT/BMP signaling overactivation, cortical hem expansion, and a shift of developmental trajectories from a forebrain towards midbrain/hindbrain identity. In contrast, perturbation during neuronal commitment (“late”) causes only modest changes without patterning defects. These findings highlight a crucial, stage-specific requirement for miRNAs in coordinating brain organoid development and tightly controlling morphogen balance for establishing regional identities.

### 5. Five forebrain-patterning miRNAs partially restore the original cell fate

To identify the specific miRNA-target interaction changes driving the molecular and cellular phenotypes in early-perturbed organoids, we selected 6 potential miRNA candidates based on the following criteria: 1) miRNAs robustly expressed during organoid development and 2) miRNAs consistently downregulated in “early” (**Fig. 6a** and **Suppl. Fig. 9a-b**).

**Fig.6.**
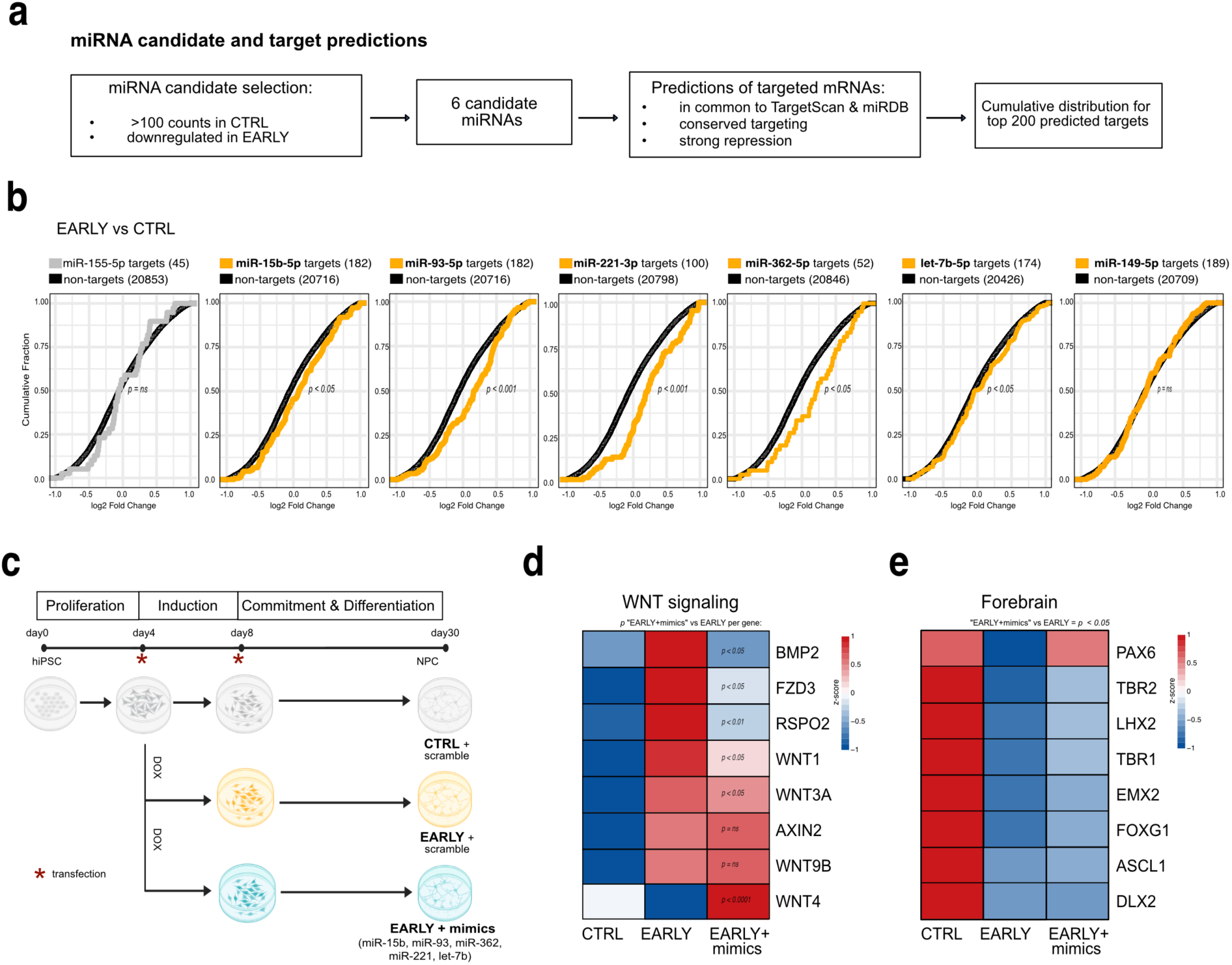
Specific pre-commitment miRNAs partially restore the forebrain specification. **a.** Pipeline for identifying miRNA candidates and their predicted mRNA targets in 30-day old organoids. miRNAs expressed in control organoids and downregulated in “EARLY” were selected. Target predictions intersected two databases (TargetScan and miRDB) and used only conserved targets with strong predicted repression. Top 200 predicted targets were used as input for downstream analyses (“Methods”). **b.** Shift of the cumulative target distribution indicates that predicted targets are preferentially upregulated upon “EARLY” perturbation, consistent with loss of miRNA activity. Cumulative distribution of log2 fold changes (EARLY vs CONTROL) for the top 200 predicted targets (yellow) versus all non-target genes (black), for each miRNA candidate. Significance determined with two-sided Kolmogorov-Smirnov test. Data from bulk RNA-seq of hiPSC line 1 (n=3 per condition; “Methods”). **c.** Design of the miRNA gain-of-function experiment (“Methods”). NPCs were transfected at two time points (asterisks) with either scramble constructs or a cocktail of five miRNA mimics in the context of early perturbation (EARLY + mimics). Cells were collected at day 30 for downstream analyses. Grey= mock control; orange= “EARLY”; cyan= “EARLY + mimics”. **d.** miRNA mimics repress the expression of some predicted WNT-pathway genes, validating those as direct targets. Normalized gene expression, quantified by qRT-PCR and z-scored across conditions in day 10 NPCs, for three replicates across three independent experiments. Significance assessed using stratified Wilcoxon-Mann-Whitney test accounting for batch effects. **e.** Forebrain transcription factor expression is partially restored by miRNA mimics. Normalized gene expression z-scored across conditions in day 30 NPCs, for three biological replicates. *PAX6, TBR2, LHX2, TBR1, ASCL1* and *DLX2* were quantified using nCounter, while *FOXG1* and *EMX1* by qRT-PCR. Global significance of the EARLY+MIMICS vs EARLY shift across all 8 genes was assessed by Wilcoxon signed-rank test and confirmed by linear mixed model.

We then examined the expression of their predicted conserved targets defined by two independent algorithms (TargetScan and miRDB, Methods). Loss of a miRNA is expected to derepress its direct targets, leading to a global trend toward target upregulation. We used the stem cell-specific miR-302 family as a positive control and miR-122 and miR-155 as negative controls (liver- and immune cell-specific miRNAs) (**Fig. 6b** and **Suppl. Fig. 9c)**. In 30-day-old early-perturbed organoids, 5 miRNA candidates, including miR-15b, miR-93, miR-221, miR-362, and let-7b, displayed a pronounced global upregulation of their predicted targets, while miR-149 did not (**Fig. 6b).** Notably, the combined enriched GO terms of the candidate miRNA predicted targets converged on WNT signaling pathways (**Suppl. Fig. 6c-d)**, supporting the idea that miRNA loss primarily derepresses WNT-related transcripts.

We next tested whether these 5 miRNAs directly contribute to the observed phenotypes. We performed a miRNA gain-of-function experiment in a neuronal-specific setup, using 2D neural forebrain progenitor cells (NPCs). We co-transfected the mimics corresponding to all 5 miRNA candidates during the pre-commitment window, concomitantly with *DROSHA* perturbation (**Fig. 6c**, Methods). First, we confirmed the upregulation of *DROSHA* in perturbed conditions (“Early” and “Early+mimics”), with a peak at day 10 (5.5-fold increase) (**Suppl. Fig. 9e**). This strong induction of *DROSHA* is in line with the activity of the neural-specific hUBC promoter. miRNA mimic expression peaked at day 10 of NPC differentiation (average of 40-fold), indicating effective miRNA availability within the appropriate temporal window, prior to lineage commitment (**Suppl. Fig. 9f**).

Predicted WNT pathway targets (*FZD3, RSPO2, WNT1,* and *WNT3A)* were significantly repressed by the mimics (**Fig. 6d** and **Suppl. Fig. 9g**), confirming WNT activation as the primary effect of the miRNA perturbation. Forebrain transcription factors previously found downregulated in early-perturbed organoids **(Fig. 4f**, **Fig. 5e),** including *PAX6* and *EMX2*, were partially restored upon mimic overexpression, to levels closer to the control conditions **(Fig. 6f** and **Suppl. Fig. 9h).**

These results demonstrate that these five miRNAs act as key pre-commitment determinants of dorsal forebrain fate, likely by constraining WNT/BMP signaling, within the pre-commitment window.

## Discussion

MicroRNAs are highly abundant in the mammalian brain, yet their precise temporal requirements in human neurodevelopment have remained elusive, largely due to the lack of suitable models.

Here, we addressed this gap by using forebrain organoids. We provide, to the best of our knowledge, the first comprehensive analysis of miRNA function in the human developing brain, showing that miRNAs are required for proper human brain development from its earliest stages. We demonstrate that our organoid model (across two independent lines) recapitulates neurodevelopmental miRNA signatures (for the cell types covered) and that neurogenic-specific miRNAs are expressed similarly to fetal forebrain, despite differences in developmental stage and cell composition (Smal et al., 2024) (**Fig. 1**). As we expected, key developmental transcriptional programs fluctuate dynamically over time (**Fig. 1c**). Interestingly, we also observed that miRNAs and their main biogenesis machinery proteins undergo temporal fluctuations in expression levels, coinciding with specific neurodevelopmental transitions (**Fig. 1d**, **Fig. 2**). This dynamic behavior is notable, given that miRNAs and their processing proteins are highly stable molecules with long half-lives (Gantier et al., 2011; Guo et al., 2015); for instance, phenotypic consequences of Dicer deletion are typically reported long after the genetic perturbation (Schaefer et al., 2007; De Pietri Tonelli et al., 2008; Kawase-Koga et al., 2009). Similar temporal expression peaks have previously been reported in simpler systems, such as Dicer during sea urchin embryogenesis (Song et al., 2011); in *C.elegans* before gastrulation (Stoeckius et al., 2009), as well as miRNA-target patterns in the human fetal brain (Nowakowski et al., 2018).

We propose that the boost in miRNA production observed at the time of cell commitment (day 15) functions to precisely regulate morphogen signaling at a critical patterning stage - when cell fates are defined and programmed (**Fig. 2**). Afterward, the reduced demand for *de novo* miRNA synthesis suggests that the established miRNA pool is sufficient to fine-tune later stage-specific processes. Whether this surge in miRNA production is exclusively required for the developmental events occurring at commitment or also primes for upcoming phases remains an open question. Notably, the uniform expression of the biogenesis machinery across cortical-like structures, including neuroepithelial loops and neurons, indicates that machinery components are ubiquitously required in different cell types (**Fig. 2h**).

Collectively, our findings support the hypothesis that, during human embryogenesis, miRNA abundance is tightly coupled to developmental stages and is not restricted to specific cell populations. This hypothesis fits with the heterochronic role identified for miRNAs in model organisms (Ruvkun et al., 1989; Lee et al., 1993; Wightman et al., 1993). To directly test miRNA temporal requirements, we globally dampened miRNA biogenesis (maturation) by expressing a dominant-negative *DROSHA* mutant at two developmental stages: prior neuronal commitment (“early,” from day 6) and during commitment (“late,” from day 10) (**Fig. 3**). This resulted in a time-dependent (pre-commitment) forebrain-midbrain imbalance (**Figs. 4-5**). We propose that, during normal human neurodevelopment, miRNA activity is required to establish forebrain identity within a specific pre-neuronal commitment window. Herein, we observed that forebrain commitment gene programs are enabled as a consequence of direct forebrain-patterning miRNA-mediated repression of WNT/BMP morphogen signaling, which consequently inhibits midbrain/hindbrain programs and in turn favors forebrain patterning (**Fig. 7A**).

**Fig.7.**
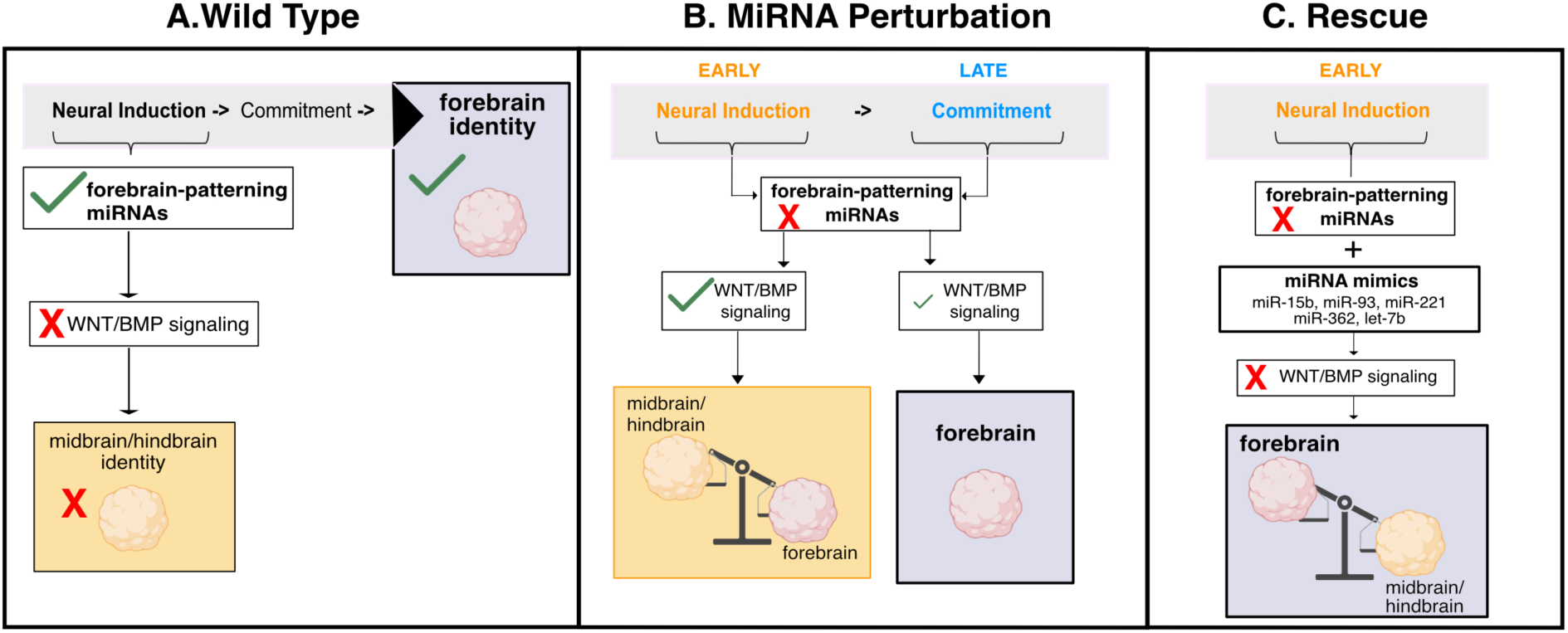
Model of miRNA temporal requirement in human neurodevelopment. **a.** We propose that miRNA activity pre-commitment is required to establish forebrain identity. Herein, forebrain patterning is enabled, while midbrain/hindbrain programs are silenced, as a consequence of direct miRNA-mediated repression of WNT/BMP morphogen signaling. **b.** When miRNAs are dysregulated during neuronal commitment (blue, right), WNT/BMP activation remains modest and the forebrain fate is preserved. In contrast, miRNA loss during neural induction (orange, right) leads to a strong derepression of WNT/BMP signaling, resulting in a patterning shift from forebrain to midbrain/hindbrain. **c.** Reintroduction of five forebrain-specific miRNAs within the pre-commitment window represses WNT/BMP signaling, consequently reversing the mis-patterning and restoring the forebrain regional identity.

When forebrain-patterning miRNAs are dysregulated during neuronal commitment, WNT/BMP activation remains mild, and the forebrain fate is preserved (**Figs. 4-5, 7B**). In contrast, forebrain-patterning miRNA loss during neural induction leads to a strong derepression of WNT/BMP signaling, resulting in a patterning shift from forebrain to midbrain/hindbrain fates (**Fig. 7B**). Notably, a similar upregulation of WNT/BMP signaling was also reported upon Drosha knockdown in sea urchin embryos (Song et al., 2011), indicating that this regulatory “heterochronic” relationship is extremely old and fundamental to tissue patterning. Reintroduction of five forebrain-patterning miRNAs during the pre-commitment window (early) represses WNT/BMP-related transcripts, consequently reversing the mis-patterning and partially restoring the forebrain regional identity (**Fig. 6-7C**). Nevertheless, we of course cannot exclude the possibility that other miRNA candidates would also influence the regionalization.

Recent work showed that temporally restricted activation (day 6 to 11) of WNT/BMP in human forebrain organoids redirects differentiation from cortical neurons toward cortical hem identities (Amin et al., 2024). Remarkably, this timeframe precisely coincides with our pre-commitment miRNA perturbation. Although we target the upstream regulators of WNT/BMP, the miRNAs, we observed a similar shift in cell fate (**Figs. 5** and **7**), indicating that timing is the critical determinant. Nevertheless, our analysis focuses on a crucial but very early and temporally restricted phase of neurodevelopment. Therefore, additional bursts in miRNA activity may occur at much later stages, regulating distinct gene programs, such as synaptogenesis, neuronal activity, axonogenesis, and synaptic pruning.

Altogether, our findings establish miRNAs as key heterochronic regulators of human neurodevelopment and delineate the specific developmental window during which their function is essential. We believe that this study will contribute to a better interpretation of the complex gene regulatory landscape of early human brain development. We also think that our study strongly argues that miRNAs should be added to the toolbox of (brain) organoid bioengineering.

## Model Limitations and Implications

Forebrain organoids faithfully model early neurogenesis but lack vascularization, microglia, and late-maturing cell types, potentially limiting insights into full cellular complexity of developing brain. Our hUBC-driven perturbation exhibited a neurogenic bias, which may understate effects in radial glia. Therefore, future Cre- or lineage-specific drivers could address this. The rescue effect was partial, suggesting contributions from additional miRNAs or indirect effects. Additionally, the rescue experiment was performed in forebrain patterned NPCs, not organoids, as efficiency of lipofection was too low in 3D models. Nevertheless, our study establishes miRNAs as essential temporal gatekeepers of human forebrain regionalization, with implications for neurodevelopmental disorders involving patterning defects (e.g., holoprosencephaly). A comprehensive understanding of how gene regulation shapes human brain development is fundamental for identifying new opportunities for disease intervention and therapeutic strategies. Together, we provide a framework for dissecting miRNA functions across brain development stages and highlight organoids as a powerful platform for such analyses.

## Author contributions

LE, CACJ, ARW, and NR conceptualized the study, with inputs from IL. LE, CACJ, ARW and NR designed the study. NR supervised LE in the project. LE, ME and ARW generated the hiPSC cell lines and derived organoids. LE, ME and JL performed the experiments. NM provided bioinformatic and experimental support. PB supported with scientific discussion and transcriptomic analyses. LE performed miRNA and bulk RNA seq bioinformatic analyses. CACJ performed all single cell RNA seq analyses. ARW and NR provided funding acquisition. LE, CACJ, ARW and NR wrote the manuscript with inputs from all authors.

## Acknowledgement

The authors thank the whole Rajewsky group and the BIMSB-MDC Organoid Platform (led by ARW) for discussions, support and feedback; Margareta Herzog for organizational help; Tancredi Massimo Pentimalli and Ilan Theurillat, for consultation about single cell RNA sequencing analysis; Svea Beier for suggestions about data visualization; Flavia Scoyni for advice about the rescue experiments and Marvin Jens for advice about the strategy to identify candidate miRNAs and target predictions. We further acknowledge Sandra Raimundo and the Microscopy Facility at MDC, for support with confocal imaging. We thank Joshua Mendell and Kenneth Chen for sharing the original plasmid containing the dominant-negative *DROSHA* variant (“V5-Drosha E1147K-pcDNA”). All illustrations were created with BioRender.com. NR thanks DFG Leibniz Award and DFG Neurocure/BrianBank for support. ARW and NR thank MDC Central Funding. LE was funded by the MDC graduate program. CCJ was funded by DFG RA 838-51 (Leibnizpreis) and HFMI Pilot Project: VirtualCell.

## Supplementary Figures

**Suppl.Fig.1.**
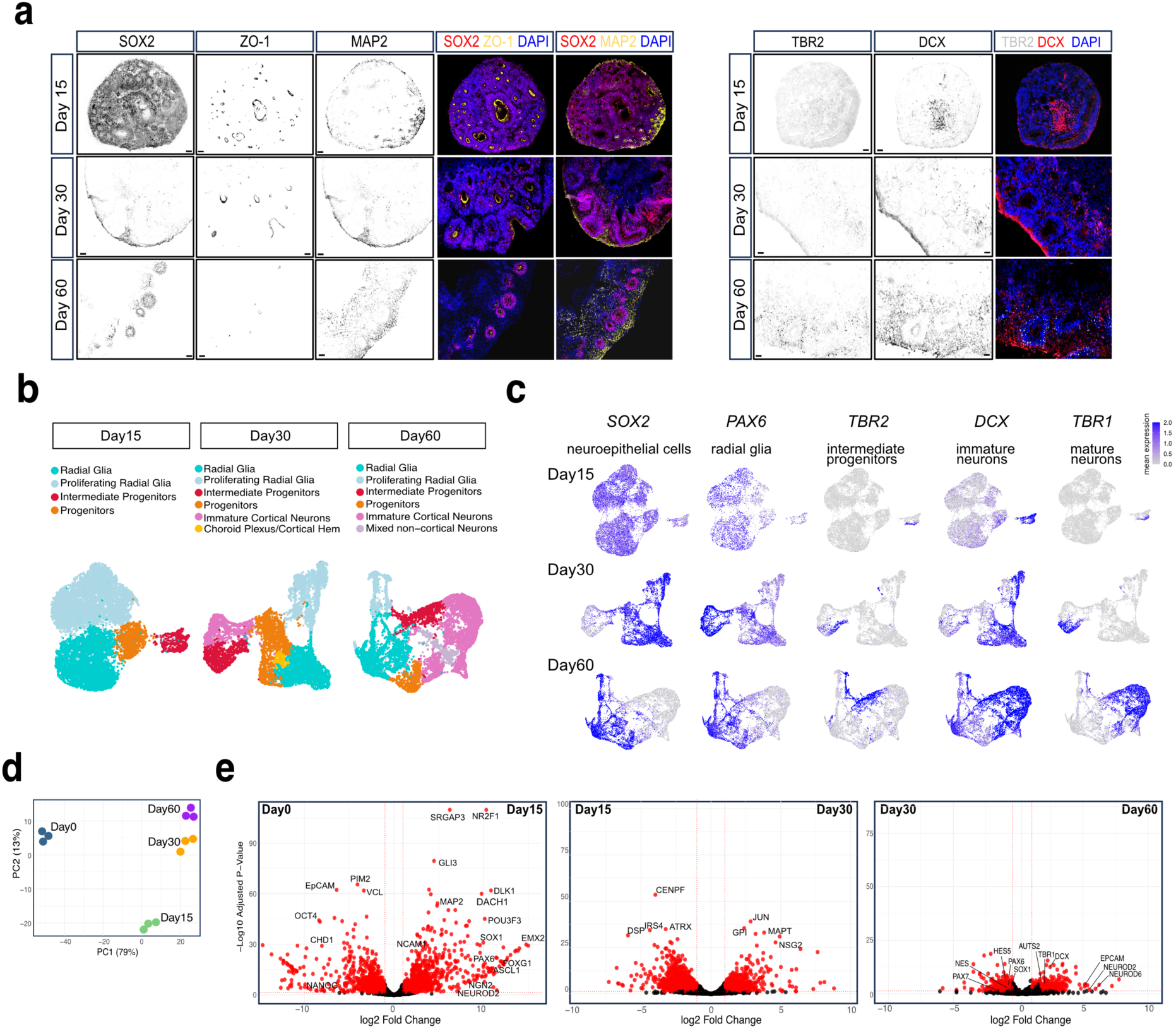
**a.** Representative immunofluorescence images of 15, 30 and 60 day-old brain organoids stained for. SOX2 (neural progenitor), ZO-1 (neuroepithelial tight junction), TBR2 (intermediate progenitors), DCX and MAP2 (neurons). Nuclei are stained by DAPI. Scale bar = 0.5 µm. **b.** UMAP of single cell RNA-sequencing analysis across organoid development. Cluster annotation at each time point is shown. Day15=pool of 5 organoids; day30 and day60=pool of 3 organoids. Data derived from hiPSC line1. **c.** Feature plots derived from single cell RNA-sequencing, displaying the distribution of exemplary markers for five main cell populations across organoid development. Day15=pool of 5 organoids; day30 and day60=pool of 3 organoids. Data derived from hiPSC line1. **d.** PCA (Principal Component Analysis) of bulk RNA-sequencing (“Methods”). Individual biological replicates (*n* = 3) are shown for hiPS cell line 1. **e.** Volcano plots showing differential gene expression analyses from bulk RNA-sequencing of hiPSCs (day0) and organoids collected at different time points (days 15,30 and 60) (“Methods”). From left to right: 1) day 15 versus day 0, 2) day 30 versus day 15, and 3) day 60 versus day 30. Red dots represent significantly differentially expressed genes (adjusted p-value > 0.05). Dotted lines indicate log2 fold change >1 or <-1. Grey: not significant. Plotted data is the mean change from three biological replicates (*n* = 3) per condition, from cell line 1.

**Suppl. Fig.2.**
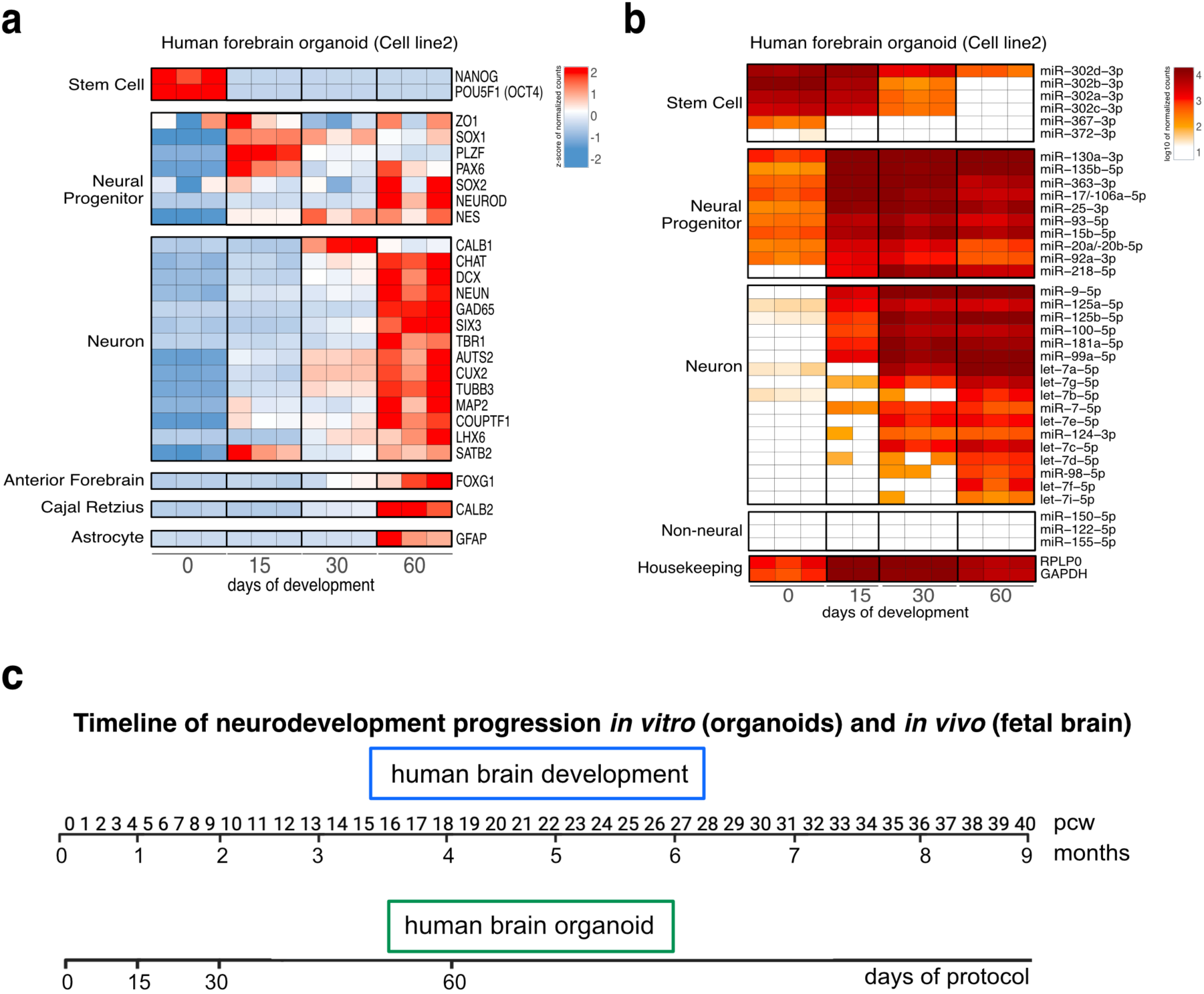
**a.** Heatmap showing the expression of selected marker genes split by cell type, over the course of organoid development. Direct quantification using nCounter (“Methods”). Expression values are displayed as z-scores of normalized counts. Each column represents a single biological replicate (Day15=pool of 6 organoids, day30 and day60= pool of 3 organoids). *n* = 3 for hiPS cell line2. **b.** Heatmap showing the expression of selected miRNAs split by cell type, over the course of organoid development (“Methods”). Direct quantification using nCounter (“Methods”). Expression values are displayed as log10 normalized counts. Each column represents a biological replicate (Day15=pool of 6 organoids, day30 and day60= pool of 3 organoids). *n* = 3 (day 0, day 30 and day 60), *n* = 2 (day15), for hiPS cell line2. **c.** Schematic showing comparison of developmental temporal progression between human brain organoids (here unpatterned, whole brain samples) and fetal brain, based on transcriptomic data from Mariani et al., 2015; Camp et al., 2015. This comparison is outlined by Kelava and Lancaster, 2016. pcw= post-conceptional week.

**Suppl. Fig.3.**
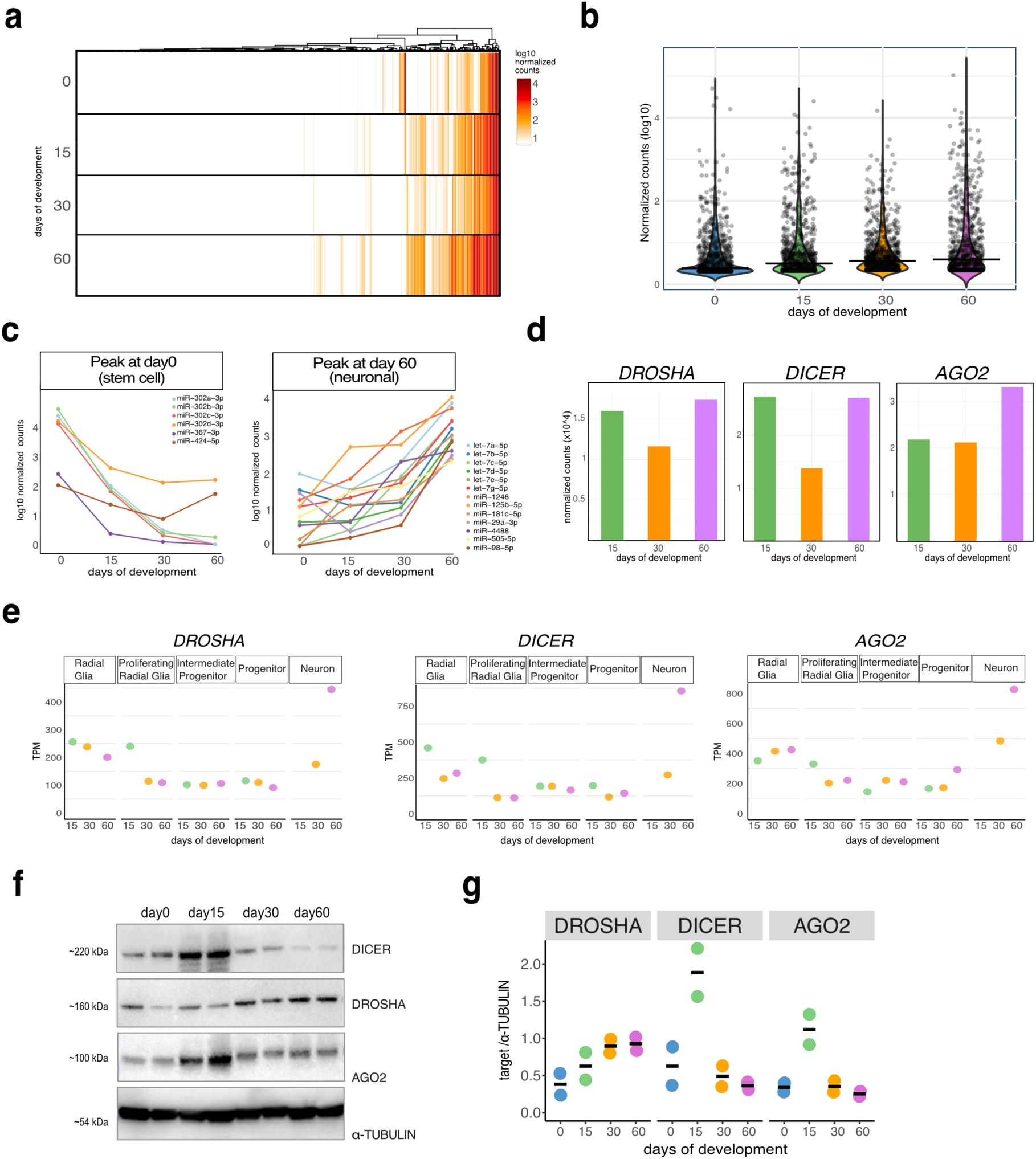
**a.** Heatmap showing the expression of microRNA species (∼800) across organoid development (“Methods”). Direct quantification using nCounter (“Methods”). Expression values are displayed as log10 normalized counts of miRNAs with no expression filtering applied. Each lane represents the mean of three biological replicates for hiPS cell line1 (Day15=pool of 6 organoids, day30 and day60= pool of 3 organoids). **b.** Violin plot showing miRNA distribution across time points, with no expression filtering applied. Direct quantification using nCounter (“Methods”). MiRNA expression is displayed as log10 normalized counts. Each dot represents the average of log2 normalized counts for *n* = 3 of hiPS cell line 1. Horizontal bar represents the median. **c.** Line plot showing expression dynamics of stem-cell specific (left, peak at day 0) and neuronal miRNAs (right, peak at day 60) over time. Direct quantification using nCounter (“Methods”). Each line represents the average of three biological replicates (Day15=pool of 6 organoids, day30 and day60= pool of 3). *n* = 3 for hiPS cell line1. **d.** Quantification of *DROSHA, DICER* and *AGO2* transcripts, based on single cell RNA-seq, analyzed as pseudobulk (all reads per sample were normalized and converted to TPM). Each bar represents one biological replicate for hiPS cell line1 (Day15=pool of 6 organoids, day30 and day60= pool of 3 organoids). TPM= transcripts per million. **e.** Quantification of *DROSHA, DICER* and *AGO2* transcripts per cell cluster, from single cell RNA-seq (Day15=pool of 6 organoids, day30 and day60= pool of 3 organoids). Each dot represents one biological replicate for hiPS cell line1 (Day15=pool of 6 organoids, day30 and day60= pool of 3 organoids). **f.** Western blot membrane showing DROSHA, DICER and AGO2 proteins over organoid development; α-TUBULIN serves as loading control. Each lane represents an individual biological replicate (Day15=pool of 8 organoids, day30 = pool of 3, “Methods”). *n* = 2 for hiPS cell line1. **g.** Western blot quantification by densitometry of DROSHA, DICER and AGO2 proteins over organoid development. Each dot represents an individual biological replicate (*n* = 2 for hiPS cell line 1). Each protein is normalized to α-TUBULIN.

**Suppl. Fig.4.**
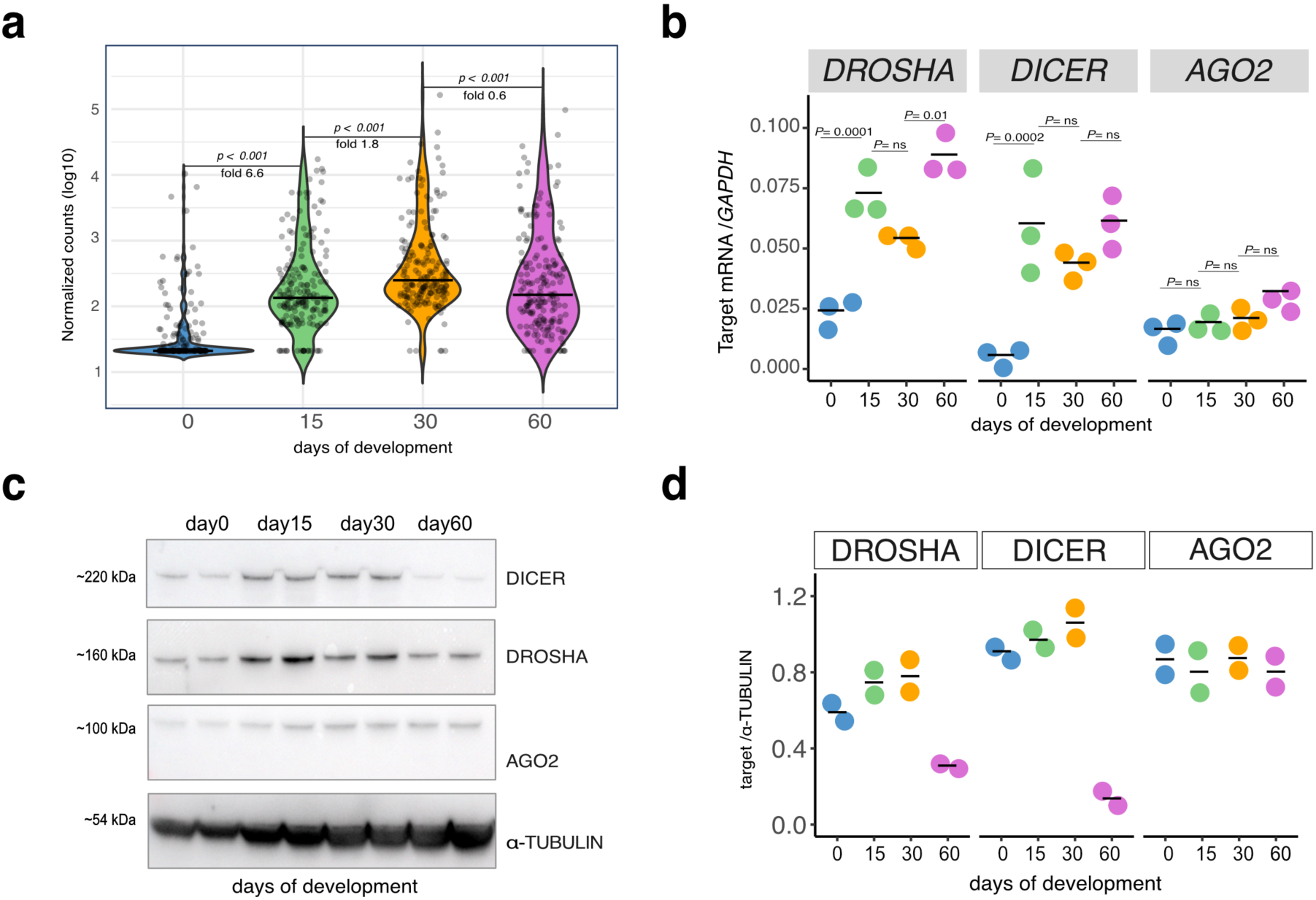
**a.** Violin plot showing miRNA distribution miRNA expression distribution over organoid development. Direct quantification using nCounter (“Methods”). Each dot represents the average log10 normalized counts (filtering: ≥ 100 counts at least at one stage) deriving from three biological replicates for days 0,30 and 60, and from two biological replicates for day15 of hiPS cell line2 (Day15=pool of 6 organoids, day30 and day60= pool of 3 organoids). Median line and fold are displayed. Significance assessed using Wilcoxon rank-sum test. **b.** qRT-PCR measurement of *DROSHA, DICER* and *AGO2* mRNAs normalized on *GAPDH*, across organoid development. Each dot represents an individual biological replicate for hiPS cell line 2 (*n* = 3). Two-sided t-test p-values computed between days, with Holm’s correction for multiple testing. **c.** Western blot membrane showing DROSHA, DICER and AGO2 proteins over organoid development; α-TUBULIN serves as loading control. Each lane represents an individual biological replicate (Day15=pool of 8 organoids, day30 = pool of 3, “Methods”). *n* = 2 for hiPS cell line 2. **d.** Western blot quantification by densitometry of DROSHA, DICER and AGO2 proteins over organoid development. Each dot represents an individual biological replicate (*n* = 2 for hiPS cell line 2). Each protein is normalized to α-TUBULIN.

**Suppl. Fig.5.**
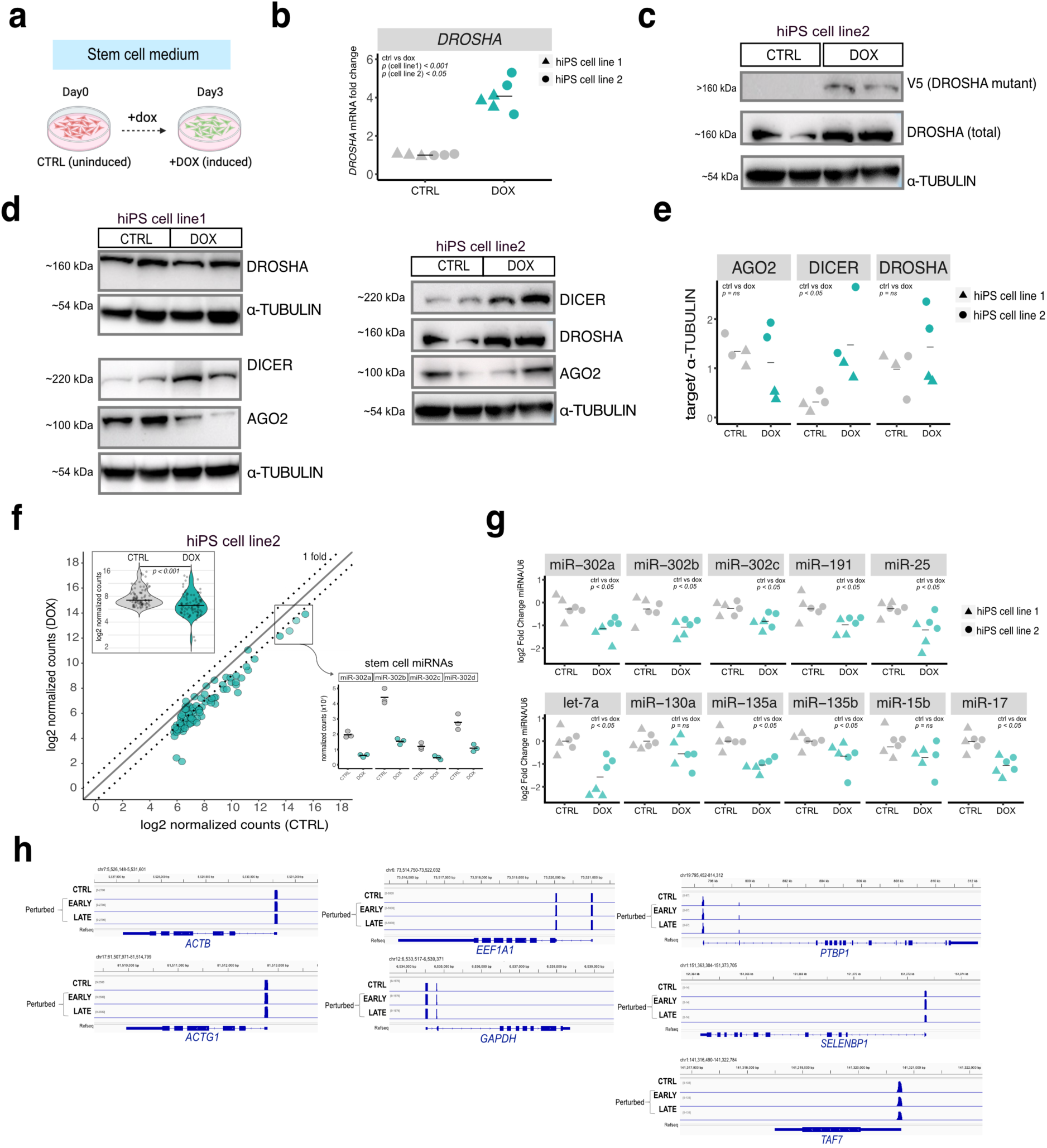
**a.** Schematic of *DROSHA* perturbation experiments conducted in hiPSCs. Cells were treated for three days with doxycycline (DOX), followed by downstream analyses. **b.** qRT-PCR measurement of total *DROSHA* mRNA. Plotted are fold changes of doxycycline (DOX)-treated over untreated control (CTRL), normalized on *GAPDH*. Each dot represents a biological replicate for two independent hiPSC lines and horizontal bars the mean of all replicates. Two-sided t-test p-values computed between control and dox-induced samples. **c.** Representative western blot membrane showing total DROSHA and V5 (DROSHA mutant) proteins in control (CTRL) and perturbed (DOX) hiPS cells (“Methods”). α-TUBULIN serves as loading control. Each lane is a biological replicate for hiPS cell line 2. **d.** Western blot membrane of miRNA biogenesis proteins in control and doxycycline (DOX)-treated cells. Each lane is a biological replicate for hiPS cell lines 1 (on the left) and 2 (on the right). α-TUBULIN serves as loading control. **e.** Western blot quantification of d. Each protein is normalized to α-TUBULIN. Horizontal bars represent the mean. For each target, significance was assessed using Wilcoxon rank-sum test, with Bonferroni correction for multiple comparisons. **f.** Scatter plot of miRNA expression in control (CTRL) and perturbed (DOX) hiPSCs, quantified by nCounter (“Methods”). Dotted lines represent a log2 fold change of ±1. Insert: violin plot showing miRNA distribution, with horizontal bar representing the median. Significance assessed using Wilcoxon rank-sum test. Each dot represents the average of log2 normalized counts (≥ 50) for three biological replicates of hiPS cell line 2. Right: quantification of stem cell-specific miRNA family (miR-302) in (CTRL) and perturbed (DOX) hiPSCs. Here, one dot represents one biological replicate for hiPS of cell line 2.. MiRNA quantification in hiPSCs done with Taqman Assays. Plotted are log2 fold changes of miRNA expression in perturbed (DOX) over control (CTRL), normalized on U6. Each dot represents an individual biological replicate (“Methods”). Data are derived from two independent cell lines. Horizontal bars represent the mean of all replicates per condition. Significance assessed using Wilcoxon rank-sum test, merging the two cell lines. P-values were adjusted for multiple testing using the Bonferroni correction. **h.** Genome browser tracks showing 5′ RNA-seq read coverage for control (CTRL) and perturbed 30-day old organoids (“EARLY” and “LATE”) of hiPS cell line1. For each condition, three organoids were pooled together (“Methods”). Peaks reflect the distribution of transcript 5′ ends, illustrating transcription start site usage.

**Suppl Fig.6.**
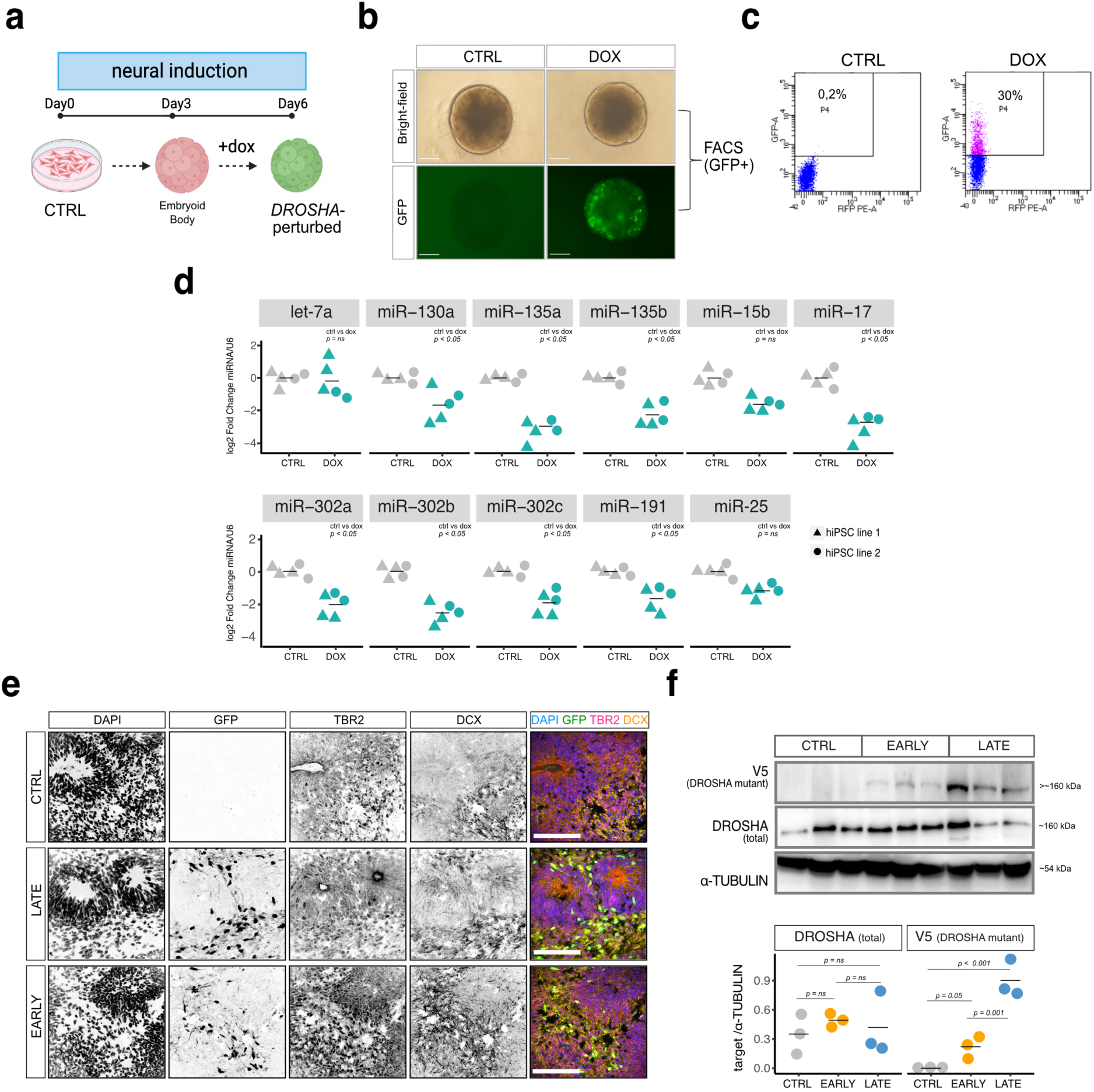
**a.** Schematic of *DROSHA* perturbation experiments in embryoid bodies (EBs). EBs were treated for three days with doxycycline (dox), before proceeding with downstream analyses. **b.** Representative microscopy images (bright-field and GFP) of control and dox-induced EBs, taken three days after doxycycline administration. Scale bar= 200um. **c.** Flow cytometry quantification plots of control and induced EBs (GFP+ cells) (“Methods”). **d.** MiRNA quantification in embryoid bodies (EBs) by Taqman Assays. Plotted are log2 fold changes of miRNA expression in GFP-sorted samples (DOX) over uninduced controls (CTRL), normalized on U6. Each dot represents an individual biological replicate (“Methods”). Data are derived from two independent cell lines. Horizontal bars represent the mean of all replicates per condition. Significance assessed using Wilcoxon rank-sum test, merging the two cell lines. P-values were adjusted for multiple testing using the Bonferroni correction. **e.** Representative immunofluorescence images (hiPS cell line1) of 30-day old control (CTRL) and perturbed (“early” and “late”) organoids stained with GFP, TBR2 (intermediate progenitors) and DCX (neurons). Nuclei are stained by DAPI. Scale bar= 50um. **f.** Top: western blot membrane showing total DROSHA and V5 (DROSHA mutant) proteins in control (CTRL) and perturbed (DOX) 30-day old organoids (“Methods”). α-TUBULIN serves as loading control. Each lane is a biological replicate for hiPS cell line 1. Bottom: western blot quantification by densitometry. Each protein is normalized to α-TUBULIN. Horizontal bars represent the mean. For each protein target, significance was assessed using Wilcoxon rank-sum test, with Bonferroni correction for multiple comparisons.

**Suppl Fig.7.**
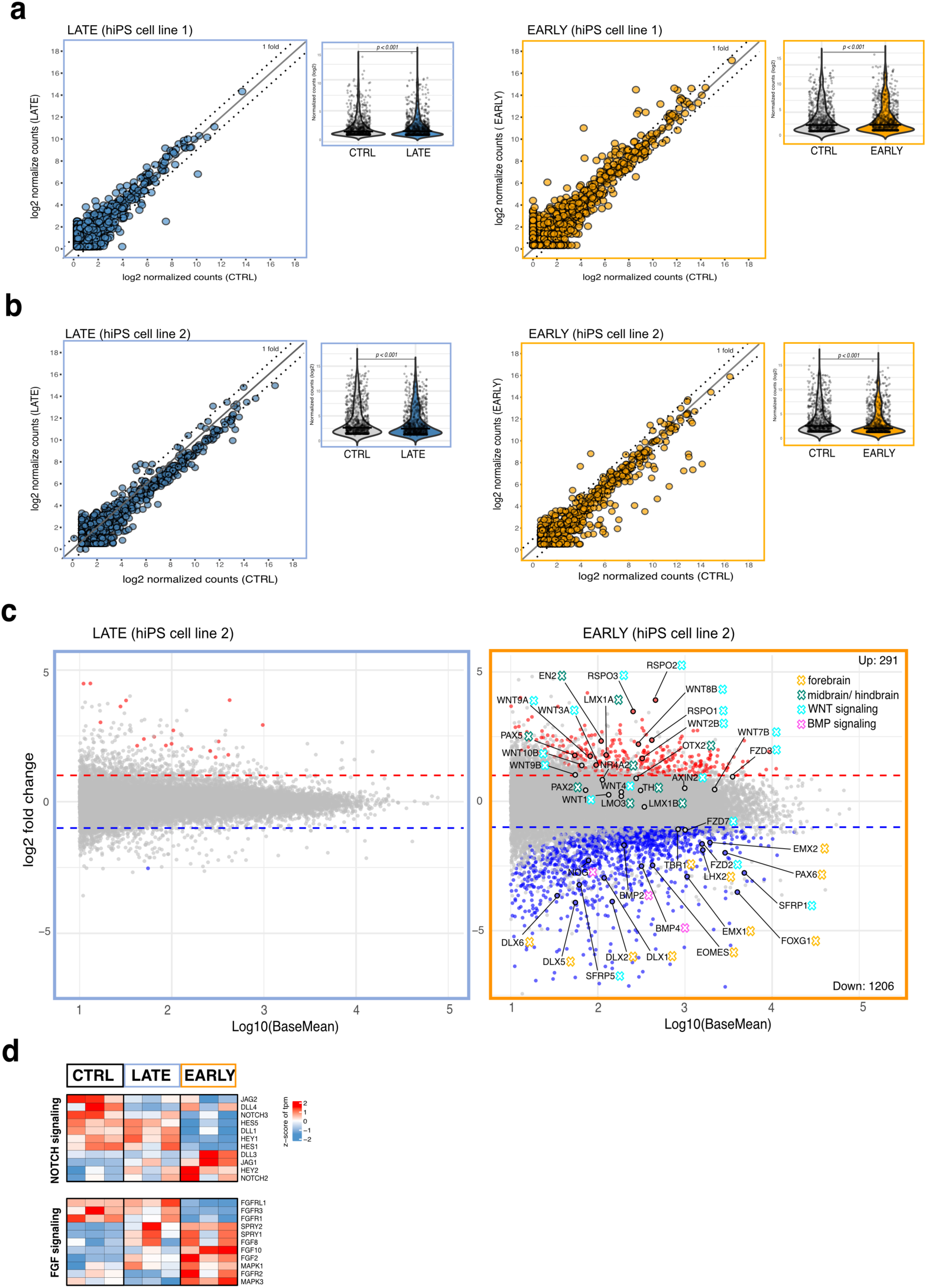
**a.** Scatter plots of miRNA expression in control (CTRL) and perturbed (“LATE” = blue; “EARLY” =orange) 30-day old organoids, quantified by nCounter (“Methods”). Dotted lines represent ±1 fold. On the right of each scatter plot: violin plot showing miRNA distribution, with horizontal bar representing the median. All miRNA species (∼800) included in the panel are displayed, with no expression filtering. Significance assessed using Wilcoxon rank-sum test. Each dot represents the average of log2 normalized counts of one miRNA, from two (left plot, “EARLY”) or three biological replicates (right plot, “LATE”) of hiPS cell line 1. **b.** Scatter plots of miRNA expression in control (CTRL) and perturbed (“LATE” = blue; “EARLY” =orange) 30-day old organoids, quantified by nCounter (“Methods”). Dotted lines represent ±1 fold. On the right of each scatter plot: violin plot showing miRNA distribution, with horizontal bar representing the median. All miRNA species (∼800) included in the panel are displayed, with no expression filtering. Significance assessed using Wilcoxon rank-sum test. Each dot represents the average of log2 normalized counts of one miRNA, for three biological replicates (*n* = 3) of hiPS cell line 2. **c.** MA plot showing differential gene expression (on the left: “LATE” vs control; on the right: “EARLY” vs control) from bulk RNA sequencing of 30-day old organoids (“Methods”). Red and blue dots respectively indicate significantly up and downregulated genes (adjusted p-value > 0.05). Dotted lines indicate log2 fold change >1 or <-1. Grey: not significant. Plotted is the mean expression change from three biological replicates (*n* = 3) per condition, from hiPS cell line 2 of two batches. One biological replicate consists of a pool of three organoids. Specific gene signatures are highlighted in colors. **d.** Heatmap displaying gene signatures for NOTCH and FGF signaling in control (CTRL), “late” and “early” 30-day old organoids, quantified with bulk RNA sequencing (“Methods”). Expression values are displayed as z-scores of tpm (transcripts per million; “Methods”). Each column represents a biological replicate (pool of 3 organoids). *n* = 3 for hiPS cell line1.

**Suppl Fig.8.**
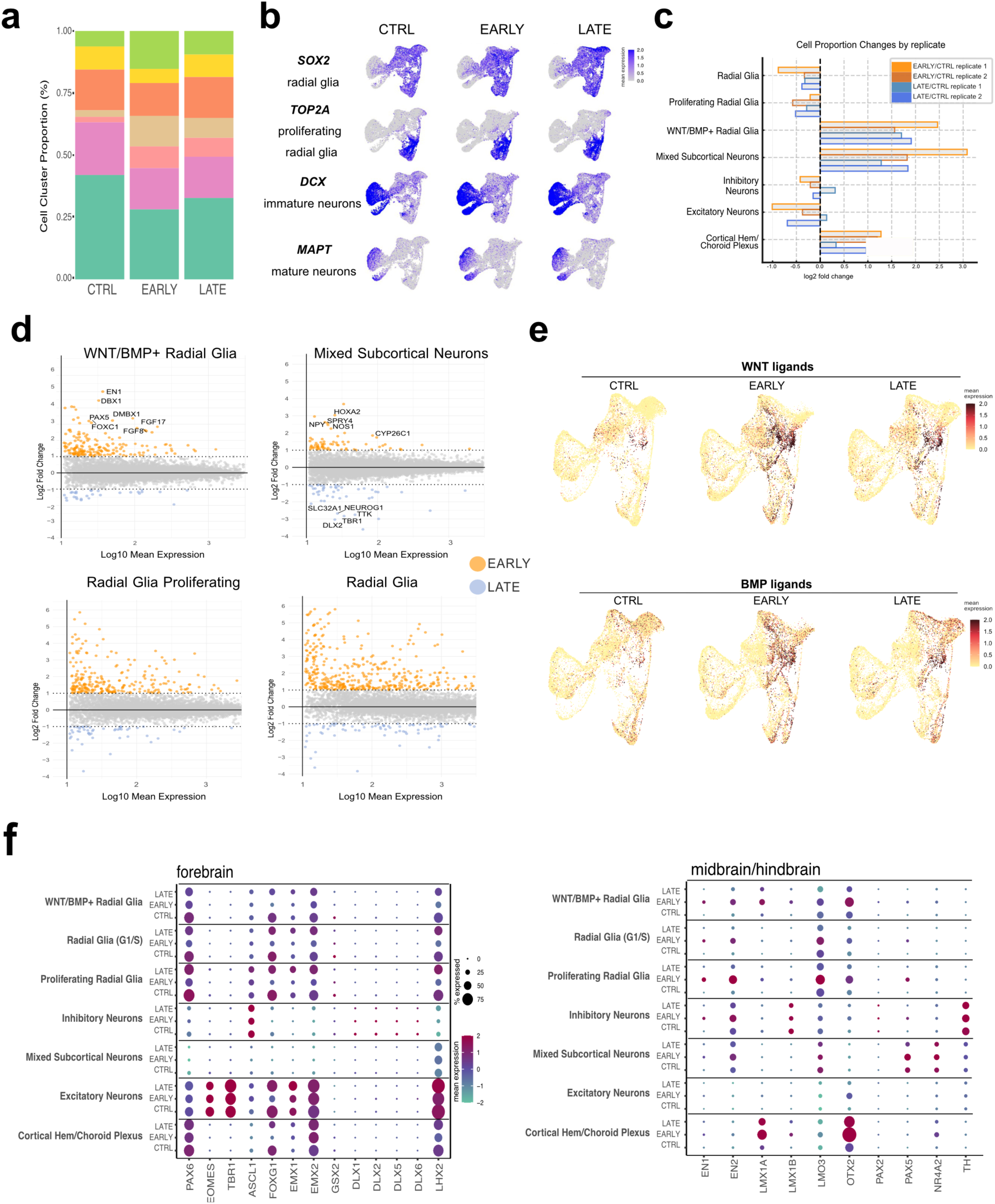
**a.** Proportions of cell types across conditions (CTRL, “EARLY”, “LATE”), from single cell RNA-sequencing analysis. Cluster annotation is shown (“Methods”). Data from two independent biological replicates per condition from hiPSC line1. **b.** Feature plots derived from single cell RNA-sequencing, displaying the distribution of exemplary markers for four main cell populations in 30-day old organoids conditions (CTRL, “EARLY”, “LATE”). Data from two independent biological replicates per condition from hiPSC line1. **c.** Proportions of cell types across conditions (CTRL, “EARLY”, “LATE”), from single cell RNA-sequencing analysis separated by replicate. Cluster annotation is shown (“Methods”). Data from two independent biological replicates per condition from hiPSC line1. **d.** MA plots showing mRNA expression changes in “EARLY” versus “LATE” for clusters of interest. Plotted is the mean change of two independent biological replicates per condition. Orange dots: “EARLY”, enriched genes (log2FC>1). Blue dots: “LATE” enriched genes (log2FC>1) (“Methods”). **e.** Uniform manifold approximation and projection (UMAP) plots of cells colored by WNT and BMP expression in each cell and separated by condition (CTRL, “EARLY”, “LATE”). Data from two independent biological replicates per condition from hiPSC line1. **f.** Dotplots showing the expression of selected marker genes in the seven major cell populations identified from single cell RNA-sequencing, across conditions (CTRL, “EARLY”, “LATE”). Separated by Forebrain (left) and midbrain/hindbrain (right). Color indicates average, scaled expression values and dot size shows the percentage of expressing cells in each condition.

**Suppl. Fig.9.**
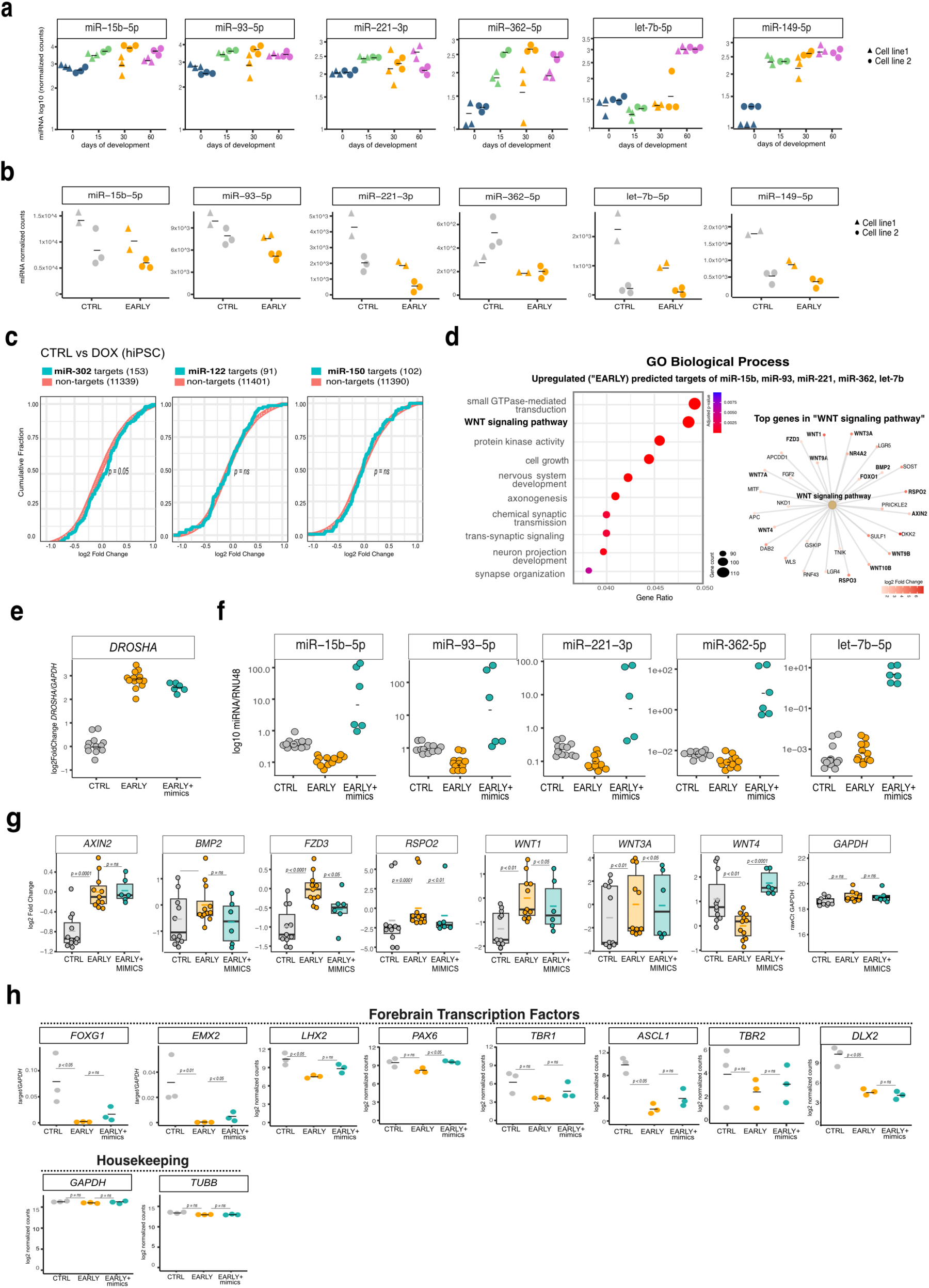
**a.** Candidate miRNA quantification over organoid development. Direct quantification using nCounter (“Methods”), expression values are displayed as log10 normalized counts of miRNAs. Each dot represents an individual biological replicate (Day15=pool of 6 organoids, day30 and day60= pool of 3 organoids). *n* = 3 for hiPS cell lines 1 and 2, and *n* = 2 for hiPS cell line2 at day15. Horizontal bars represent the mean of all replicates per time point. **b.** Candidate miRNA quantification in 15-day old organoids. Direct quantification using nCounter (“Methods”), expression values are displayed as log10 normalized counts of miRNAs in CTRL and “EARLY” conditions. Each dot represents an individual biological replicate (Day15=pool of 6 organoids, day30 and day60= pool of 3 organoids). n = 3 for hiPS cell line 2, and n = 2 for hiPS cell line 1. **c.** Cumulative distribution function (CDF) plot of gene expression comparing top 200 predicted targets of miRNA candidates (in cyan) to non-target genes (in red) in hiPSCs. Log2 fold changes derived from differential gene expression analysis of bulk RNA-seq of control (CTRL) versus perturbed hiPS of cell line 1 (n=3) (“Methods”). Significance determined with two-sided Kolmogorov-Smirnov test. **d.** Gene Set Enrichment Analysis (GSEA) on Gene Ontology (GO) BP (Biological Process) term, performed on the combined, upregulated (“EARLY” vs control) predicted targets for the 5 miRNA candidates over background genes. Left: the top 10 enriched GO terms are displayed. P-values: Benjamini-Hochberg-corrected GSEA test. The size of each dot corresponds to the number of genes associated with the respective GO term and the color gradient to the enrichment significance, expressed as adjusted p-value. Right: network plot showing the association between the combined, upregulated (“EARLY” vs control) predicted targets for the five miRNA candidates and enriched Gene Ontology (GO) terms. The central beige nodes represent significantly enriched GO terms, while the outer the individual genes associated with these terms. The color intensity and the size of each gene node correspond to the magnitude of the log₂ fold change, with larger darker nodes indicating higher upregulation. Edges indicate gene-to-GO-term associations. **e.** qRT-PCR measurement of total *DROSHA* mRNA in NPCs collected at day 10 of protocol. Plotted are log2 fold changes of 1) control (CTRL) over “EARLY” and 2) “EARLY+mimics” over “EARLY”, normalized on *GAPDH*. Each dot represents a biological replicate from three independent experiments conducted in NPCs. Horizontal lines denote the mean. **f.** MiRNA quantification in NPCs collected at day 10 of protocol by Taqman Assays. Plotted is log10 miRNA expression relative to RNU48. Each dot represents an individual biological replicate (“Methods”). Each dot represents a biological replicate from three independent experiments conducted in NPCs. Horizontal lines denote the mean. **g.** miRNA mimics repress the expression of some predicted WNT-pathway genes, validating those as direct targets. qRT-PCR was performed in NPCs collected at day 10. Log2 fold changes are shown for control (CTRL) versus “EARLY” and for “EARLY+mimics” versus “EARLY”, normalized on *GAPDH*. Raw *GAPDH* Cts values serve as technical control. Each dot represents one replicate across three independent experiments. Horizontal lines within each box denote the median. Significance assessed using stratified Wilcoxon-Mann-Whitney test accounting for batch effects. **h.** Quantification of forebrain mRNAs across conditions in NPCs collected at day 30 of protocol (“Methods”). Direct quantification using either qRT-PCR (for *FOXG1* and *EMX1*) or nCounter. Expression values are displayed as expression relative to *GAPDH* for qRT-PCR or as normalized counts for nCounter (“Methods”). Each dot represents one biological replicate.

